# Conserved accessory genes link a phylogenetically distinct *Bacillus subtilis* strain from Indian bekang to the Japanese natto clade

**DOI:** 10.1101/2025.07.08.663414

**Authors:** Kiyohiko Seki, Yukio Nagano

## Abstract

*Bacillus subtilis* is central to Asian fermented soybean foods, including Japanese natto. To explore the genomic boundaries of *B. subtilis* var. *natto*, we conducted a comparative pangenome analysis of 42 strains, including the core natto clade (n=26) and its closest relatives. Our analysis revealed a striking evolutionary paradox centered on a single strain isolated from Indian bekang. Core-genome phylogenetic analysis places this bekang strain clearly outside the tight natto clade, with a Nepalese kinema strainbeing its closest systematic neighbor. In stark contrast, quantitative analysis of accessory gene profiles revealed this single bekang strain is the functional nearest neighbor to the natto clade, sharing a highly conserved accessory gene repertoire. This shared profile defines a “natto-type” adaptive strategy (the “broad-sense natto group,” n=27), separating it from other related strains. Analysis of this group-specific repertoire revealed an enrichment of transcriptional regulators and metabolic enzymes. This finding provides a compelling case study (n=1) of polygenic adaptation, suggesting complex evolutionary pathways, such as horizontal gene transfer or selective retention, can drive rapid adaptation across disparate lineages.

## Introduction

Soybeans are a globally important protein source and have been a primary ingredient in a diverse array of Asian fermented foods for centuries, including staples like soy sauce, miso, tempeh, fŭrŭ, and douchi^1^. Fermentation not only boosts the nutritional value and shelf life of soybeans but also imparts unique flavors and functionalities, profoundly enriching regional food cultures. Central to the production of many of these soybean products is *Bacillus subtilis*^2^. This bacterium plays a pivotal role in creating foods like Japanese natto, bekang from India and Myanmar, Korean cheonggukjang, and Nepalese kinema^2^. In these foods, *B. subtilis* is crucial for breaking down soy proteins and carbohydrates, thereby generating umami compounds, distinct aromas, and, in some cases, viscous substances such as polyglutamic acid (PGA)^2,3^.

The properties of these foods, and the *B. subtilis* strains that produce them, vary significantly. Strains used for Japanese natto (produced by *B. subtilis* var. *natto*) and Korean cheonggukjang are extensively researched^4,5^. Phenotypically, natto is characterized by its intense, PGA-derived viscosity, whereas cheonggukjang is generally much less viscous. In contrast, other regional foods like Indian bekang are far less documented^6,7^. Bekang is a traditional food consumed in Mizoram, India, and adjacent areas. Critically, despite the scarcity of research, a report indicates that bekang, much like natto, is also characterized by significant viscosity and stringiness, a trait linked to PGA production^7^.

This striking convergence of a key phenotype (viscosity) in geographically distant foods (Japanese natto and Indian bekang) makes a genomic comparison particularly compelling. Our research, therefore, focuses on the industrially and culturally significant *B. subtilis* var. *natto* clade. A fundamental question remains: What defines the genomic boundary of this important industrial clade? That is, what genomic features (particularly in the accessory genome) distinguish the natto clade from its phylogenetically closest relatives? Understanding this boundary is key to unraveling the evolutionary mechanisms that led to this specific niche adaptation. Recent advances in genomics now provide access to sequence information for many *B. subtilis* strains, enabling this focused comparison^6,8,9,10,11,12,13,14,15,16,17,18,19,20,21,22,23,24,25,26,27,28^.

To unravel these genetic relationships, pangenome analysis offers a powerful framework^29,30,31^. This method dissects the collective genome of a microbial population into ‘core genes’ (reflecting vertical evolution and phylogeny) and ‘accessory genes’ (reflecting niche adaptation). While broad pangenome studies of the *B. subtilis* species group exist^32,33^, their large scale is not optimized for resolving the fine-grained genomic differences between a specific industrial clade (like *B. subtilis* var. *natto*) and its immediate sister groups.

This distinction raises a fundamental question, now applied with greater focus: Do the phylogenetic relationships inferred from core genes necessarily align with the accessory gene profiles that reflect adaptation to the nutritional niche of soybean fermentation? This refers to the specific selective pressures of an environment characterized by high concentrations of soy proteins and carbohydrates, requiring efficient enzymatic degradation.

Therefore, the objective of this study was to specifically target the *B. subtilis* var. *natto* clade and its closest relatives to address this question. Our goal was to compare the systematic boundary of the natto clade (defined by core-genome phylogeny) with its functional boundary (defined by accessory-gene profiles). We sought to determine if the phylogenetically nearest neighbor to the natto clade was also its functionally nearest neighbor, or if a discrepancy existed, which would suggest complex evolutionary pathways such as polygenic adaptation.

In pursuing this objective, we made a remarkable discovery centered on a single *B. subtilis* strain isolated from Indian bekang. We report here this compelling case study, which presents a genomic paradox: this single strain, while clearly phylogenetically distinct from the *B. subtilis* var. *natto* clade, exhibits a striking functional similarity based on a shared accessory gene repertoire. As we will demonstrate, this finding provides a key insight into the complex evolutionary pathways of microbial niche adaptation.

## Results

### Initial Genome Collection and Overall Pangenome Structure

To explore the genomic diversity surrounding the *B. subtilis* var. *natto* lineage, we initially assembled a dataset of 55 *B. subtilis* genomes, selected based on DNA sequence similarity searches using *B. subtilis* var. *natto* sequences as queries and relevant literature. This initial collection is detailed in Table 1. The set includes numerous strains from Asian fermented food production, alongside several environmental isolates, intended for an exploratory analysis to identify a focal group for deeper investigation. All genomes met high-quality standards for completeness and contamination (Supplementary Table 1). Exploratory pangenome analysis of the 55 strains revealed a large and open pangenome (Supplementary Figure 1). The pangenome size consistently increased with the addition of new genomes, indicating significant genetic diversity. Conversely, the core genome stabilized at approximately 3,000 genes.

**Table 1.** Details of the publicly available data used in this study.

| Name | Group<sup>a</sup> | Accession number | Strain name | Origin details | Location | Publication |
| --- | --- | --- | --- | --- | --- | --- |
| Clin_JPN_1 | N | GCF_037094835.1 | TUM22925 | Patient blood | Japan | 8 |
| Clin_JPN_2 | N | GCF_037094855.1 | TUM22926 | Patient bronchoalveolar lavage fluid | Japan | 8 |
| Clin_JPN_3 | N | GCF_037095795.1 | TUM22927 | The natto with the patient usually eating | Japan | 8 |
| Coll_1 | N | GCF_002982175.1 | ATCC 21228 | N/A | N/A | N/A |
| Coll_2 | N | GCF_024508195.1 | BGSC 27E1 | N/A | N/A | N/A |
| Coll_3 | N | GCF_001565875.1 | CGMCC 2108 | Soil | N/A | 9 |
| Coll_4 | N | GCF_024507815.1 | BGSC 27E3 | N/A | N/A | N/A |
| Env_CHN_1 | N | GCF_002096095.1 | Bs-115 | Soil of a corn field | Shandong, China | 10 |
| Env_CHN_2 | N | GCF_045343855.1 | G01 | N/A | Guangdong, China | N/A |
| Env_CHN_3 | N | GCF_001890405.1 | KH2 | Soil | Tianjin, China | 11 |
| Env_CHN_4 | N | GCF_023278545.1 | ZHX3 | Activated sludge | Changsha, China | N/A |
| Env_JPN_1 | N | GCF_001902555.1 | 29R7-12 | Coal Bed | Shimokita Peninsula, Japan | 12 |
| Food_CHN_1 | N | GCF_022701095.1 | BN0 | Soybean product | Wuxi, China | 13 |
| Food_CHN_2 | N | GCF_020773035.1 | SCP010-1 | Bean Paste | Sichuan, China | N/A |
| Food_IDN_1 | N | GCF_027923485.1 | G8 | Natto | Indonesia | 14 |
| Food_JPN_1 | N | GCF_003665335.1 | N1-1 | Natto | Japan | N/A |
| Food_JPN_2 | N | GCF_003665315.1 | N2-2 | Natto | Japan | N/A |
| Food_JPN_3 | N | GCF_003665355.1 | N3-1 | Natto | Japan | N/A |
| Food_JPN_4 | N | GCF_033412715.1 | N3378-2at | Natto | Hokkaido, Japan | 15 |
| Food_JPN_5 | N | GCF_033413615.1 | N3378-3At | Natto | Hokkaido, Japan | 15 |
| Food_JPN_6 | N | GCF_003665295.1 | N4-2 | Natto | Japan | N/A |
| Food_JPN_7 | N | GCF_000209795.2 | BEST195 | Natto | Japan | 16 |
| Food_JPN_8 | N | GCF_030297075.1 | NARUSE | Natto | Japan | 17 |
| Food_JPN_9 | N | Assembled from DRR029699* | TAKAHASHI | Natto | Japan | 18 |
| Food_KOR_1 | N | GCF_014961985.1 | CMIN-4 | Cheonggukjang | Jeonbuk, South Korea | N/A |
| Food_KOR_2 | N | GCF_004323065.1 | VK161 | Natto | Seongnam, South Korea | 19 |
| Food_IND_1 | Bekang | GCF_018986855.1 | Miz-8 | Bekang | Mizoram, India | 6 |
| Food_IND_2 | R | GCF_037169145.1 | Tamang | Kinema | Sikkim, India | 20 |
| Food_KOR_3 | R | GCF_003665395.1 | 2KL1 | Cheonggukjang | Jeonju, South Korea | N/A |
| Food_KOR_4 | R | GCF_003665275.1 | 2RL2-3 | Cheonggukjang | Gyeonggi, South Korea | N/A |
| Food_KOR_5 | R | Assembled from DRR029692* | NFRI8338 | Cheonggukjang | Seoul, South Korea | 18 |
| Food_KOR_6 | R | GCF_002269175.1 | DKU_NT_02 | Cheonggukjang | Cheonan, South Korea | 21 |
| Food_KOR_7 | R | GCF_002269195.1 | DKU_NT_03 | Cheonggukjang | Cheonan, South Korea | 22 |
| Food_KOR_8 | R | GCF_003665195.1 | GFR-12 | Cheonggukjang | Gyeonggi, South Korea | N/A |
| Food_KOR_9 | R | GCF_001747445.1 | HRBS-10TDI13 | Doenjang | Sunchang, South Korea | N/A |
| Food_KOR_10 | R | GCF_033113625.1 | J46 | Doenjang | South Korea | 23 |
| Food_KOR_11 | R | GCF_046840395.1 | KFRI-P74 | Fermented soybean paste | Ansung, South Korea | N/A |
| Food_KOR_12 | R | GCF_003665235.1 | MH-1 | Cheonggukjang | Chungcheongbuk-do, South Korea | N/A |
| Food_KOR_13 | R | GCF_035621695.1 | PN1236 | Fermented soybean food | South Korea | N/A |
| Food_MMR_1 | R | Assembled from DRR029694* | NFRI8316 | Pe poke | Yangon, Myanmar | 18 |
| Food_NPL_1 | R | Assembled from DRR029696* | NFRI8292 | Kinema | Hille, Nepal | 18 |
| Food_THA_1 | R | Assembled from DRR029695* | NFRI8347 | Thua nao | Chiang Mai, Thailand | 18 |
| Coll_5 |  | GCF_000009045.1 | 168 | N/A | N/A | 24 |
| Coll_6 |  | GCF_004101945.1 | ATCC 11774 | N/A | N/A | N/A |
| Env_DZA_1 |  | GCF_002142595.1 | TLO3 | Olive tree rhizosphere | Tlemcen, Algeria | 25 |
| Env_EGY_1 |  | GCF_009497795.1 | MENO2 | Honeybee gut | Egypt | 26 |
| Env_IND_1 |  | GCF_018454405.1 | A1 - Midalam | Beach soil | Tamil Nadu, India | 27 |
| Env_USA_1 |  | GCF_037976695.1 | FSL K6-1366 | HTST Pasteurization | N/A | N/A |
| Env_USA_2 |  | GCF_047302535.1 | SY1483 | Root | USA | N/A |
| Food_BFA_1 |  | GCA_029536855.2 | PRO120 | Maari | Toulfe, Burkina Faso | N/A |
| Food_IND_3 |  | GCF_018224865.2 | SC3.7 | Soybean seed Chutney | Manipur, India | 28 |
| Food_KOR_14 |  | GCF_030285965.1 | KFRI-FA111 | Jeotgal | South Korea | N/A |
| Food_KOR_15 |  | GCF_003665215.1 | PJ-7 | Cheonggukjang | Gyeonggi, South Korea | N/A |
| Food_KOR_16 |  | GCF_009913275.1 | SRCM102754 | Doenjang | South Korea | N/A |
| Food_LAO_1 |  | Assembled from DRR029693* | NFRI8302 | Thua nao | Ruanpanpan, Laos | 18 |

\*Assembled *de novo* from Sequence Read Archive data. Abbreviations used in "Group": N = "Narrow-sense natto group" (n=26), R = "Natto-related group" (n=15), Bekang = the single Bekang isolate (Food\_IND\_1). Abbreviations used in "Name": Clin – Clinical study; Coll – Collection; Env – Environmental; JPN – Japan; CHN – China; IDN – Indonesia; IND – India; KOR – South Korea; MMR – Myanmar; NPL – Nepal; THA – Thailand; DZA – Algeria; EGY – Egypt; BFA – Burkina Faso; LAO – Laos.

### Focusing the Analysis on the Natto Clade and Its Relatives

Our primary interest lies in the industrially important natto strains. Exploratory phylogenetic (core-genome tree) and functional (accessory-gene cluster, based on Jaccard distance) analyses of all 55 strains (Supplementary Figures 2, 3, and 4) were conducted. However, neither analysis showed clear, distinct boundaries separating all strains into discrete groups. As the primary objective of this study is to compare the *B. subtilis* var. *natto* clade with its closest relatives, we focused our main analysis on a subset of these strains. We first identified a highly homogeneous, tight-knit clade (n=26) containing all *B. subtilis* var. *natto* strains, which we hereafter refer to as the “narrow-sense natto group”. To create a relevant comparison set, we selected an additional 16 strains (for a total of 42) that were relatively close to the natto clade in both exploratory analyses and were also isolated from fermented soybean foods. The remaining 13 strains were excluded from the main analysis as they were either phylogenetically or functionally distant from the natto clade, and this set also included a relatively high number of environmental isolates.

### Core-Genome Phylogeny Reveals a Systematic “Nearest Neighbor”

We constructed a core-genome phylogenetic tree for the 42 strains (Figure 1, Supplementary Figure 5). This tree clearly separates the “narrow-sense natto group” (n=26) from the other 16 related strains. Notably, within this related set, the strain isolated from Nepalese kinema (Food_NPL_1) is positioned as the systematic nearest neighbor (the most recent outgroup) to the “narrow-sense natto group” clade. The Indian bekang isolate (Food_IND_1) is placed more distantly within this related group, clearly separated from the “narrow-sense natto group”.

**Figure 1.**
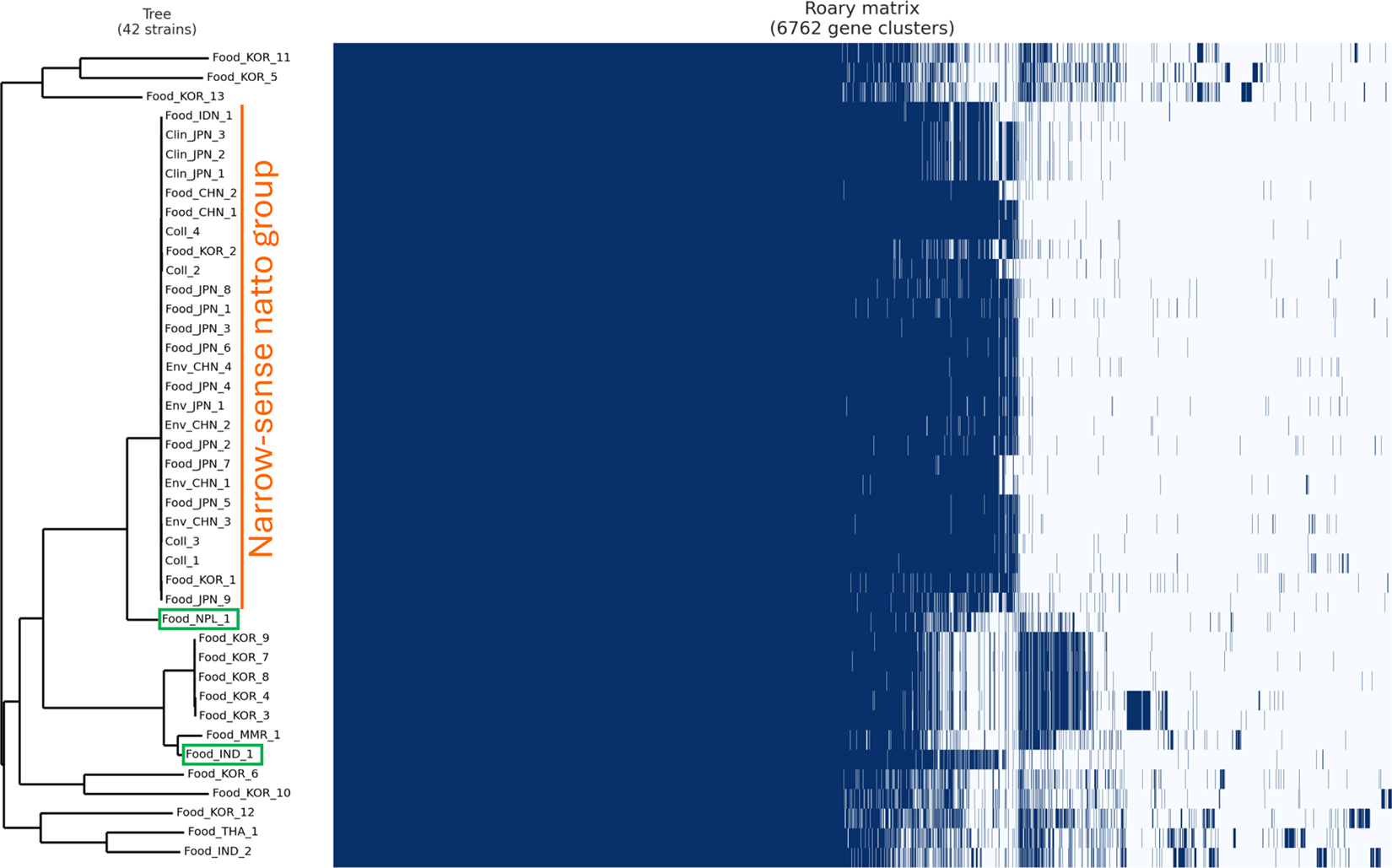
Core-genome-based phylogeny and accessory gene presence/absence matrix of the 42 *B. subtilis* strains. The phylogenetic tree on the left was constructed from a core genome alignment of 42 strains using the maximum likelihood method and is presented as midpoint-rooted. The matrix on the right, generated by Roary, visualizes the pangenome, which consists of 6,762 gene clusters. Each row corresponds to a strain in the tree, and each column represents a single gene cluster. Dark blue cells indicate the presence of a gene cluster in a strain, while white cells indicate its absence. The “narrow-sense natto group” (n=26) is indicated by an orange line. Key strains discussed in the text, the systematic nearest neighbor (Food_NPL_1) and the functional nearest neighbor (Food_IND_1), are highlighted with green squares on their labels.

### Accessory Genome Clustering Reveals a Contradictory “Functional Nearest Neighbor”

Intriguingly, even a visual inspection of the gene presence/absence matrix associated with the 42-strain core-genome tree (Figure 1, Supplementary Figure 5) provides an intuitive hint that the bekang strain (Food_IND_1), despite its phylogenetic position, shares a profile somewhat similar to the narrow-sense natto group. To formally and quantitatively assess these functional relationships, we performed hierarchical clustering on the 42 strains based on their accessory gene presence/absence profiles using Jaccard distance as the metric (Figure 2). This quantitative analysis revealed a striking paradox. The resulting cluster map (Figure 2) shows a large, functionally similar cluster corresponding to the “narrow-sense natto group” (n=25). However, the functional nearest neighbor (the most recent outgroup) to this cluster is not the systematic nearest neighbor (Food_NPL_1). Instead, it is the Indian bekang isolate (Food_IND_1). The bekang strain, which is phylogenetically more distant (Figure 1, Supplementary Figure 5), is functionally more similar to the “narrow-sense natto group” than any other strain in this set. The systematic nearest neighbor (Food_NPL_1), despite its close phylogenetic relationship (Figure 1), clusters immediately outside the functional group formed by the “narrow-sense natto group” and the bekang strain in the accessory gene analysis (Figure 2).This clear “discrepancy” — where the systematic nearest neighbor (Food_NPL_1) and the functional nearest neighbor (Food_IND_1) are not the same strain — forms the central finding of this study. This quantitative analysis was used as an exploratory map to assess functional similarity, rather than to define statistically discrete groups.

**Figure 2.**
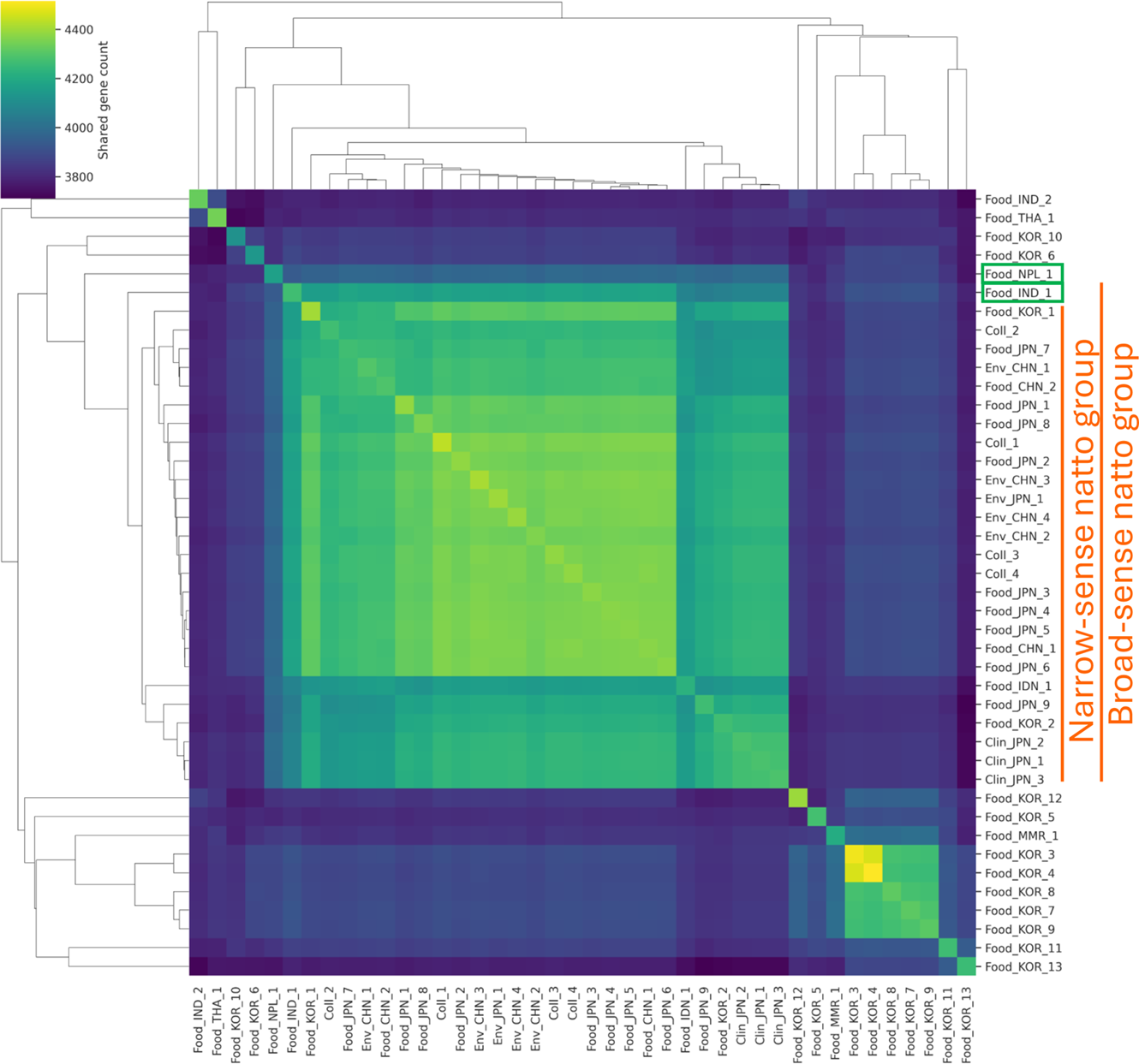
Hierarchical clustering of 42 *B. subtilis* strains based on accessory gene profiles. The heatmap shows the pairwise shared gene counts between all 42 strains. Hierarchical clustering was performed using Jaccard distance calculated from the binary presence/absence matrix. The “narrow-sense natto group” (n=26) and the “broad-sense natto group” (n=27) are indicated by orange lines. Key strains discussed in the text, the systematic nearest neighbor (Food_NPL_1) and the functional nearest neighbor (Food_IND_1), are highlighted with green squares on their labels.

### Synteny and Phylogenetic Network Analyses

To investigate the genomic basis for the functional similarity despite phylogenetic distance, we performed a chromosome-wide synteny analysis comparing representative strains with available chromosome-level genome sequences (Figure 3, Supplementary Figure 6). We selected the natto strain Food_JPN_7 (“narrow-sense natto group”), the bekang strain Food_IND_1, and the doenjang strain Food_KOR_9 (“natto-related group”). The results revealed relatively fewer insertions and deletions (indels) between Food_JPN_7 and Food_IND_1, compared to the significantly larger number of indels between the Food_JPN_7/Food_IND_1 pair and Food_KOR_9. A similar pattern of fewer indels between the natto/bekang pair compared to the bekang/related-group pair was observed when using another available chromosome-level genome from the “natto-related group,” Food_KOR_11 (Supplementary Figure 6).

**Figure 3.**
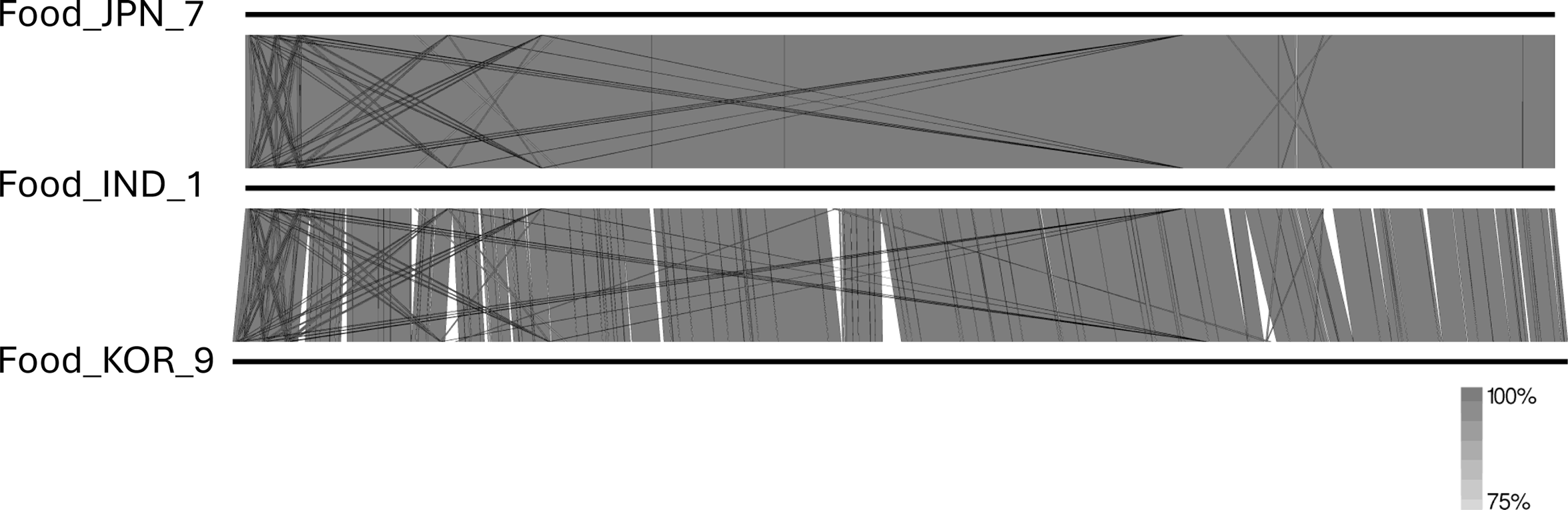
Chromosome-wide synteny analysis of representative strains. This visualization compares the complete chromosome sequences of three strains, represented by horizontal bars: *B. subtilis* var. *natto* BEST195 (Food_JPN_7, top bar), the bekang isolate (Food_IND_1, middle bar), and *B. subtilis* HRBS-10TDI13 (Food_KOR_9, bottom bar). The shaded regions connecting the genomes denote areas of nucleotide similarity identified by BLASTn, with the grayscale intensity corresponding to the percentage identity shown in the scale bar. The top panel shows the pairwise comparison between the natto strain (Food_JPN_7) and the bekang isolate (Food_IND_1). The bottom panel shows the pairwise comparison between the bekang isolate (Food_IND_1) and the “natto-related group” strain (Food_KOR_9).

Next, we investigated whether recombination could explain the similarity. We constructed a phylogenetic network from the core genome alignment for the 42 strains (Figure 4, Supplementary Figure 7). We did not detect a clear reticulate signal connecting the bekang isolate (Food_IND_1) directly to the “narrow-sense natto group”. However, the analyses did reveal complex relationships among other strains, suggesting recombination and HGT have occurred within this population. Of particular interest was evidence of recombination between the bekang isolate and the *B. subtilis* strain (Food_MMR_1) from Myanmar, regions which share a geographical border.

**Figure 4.**
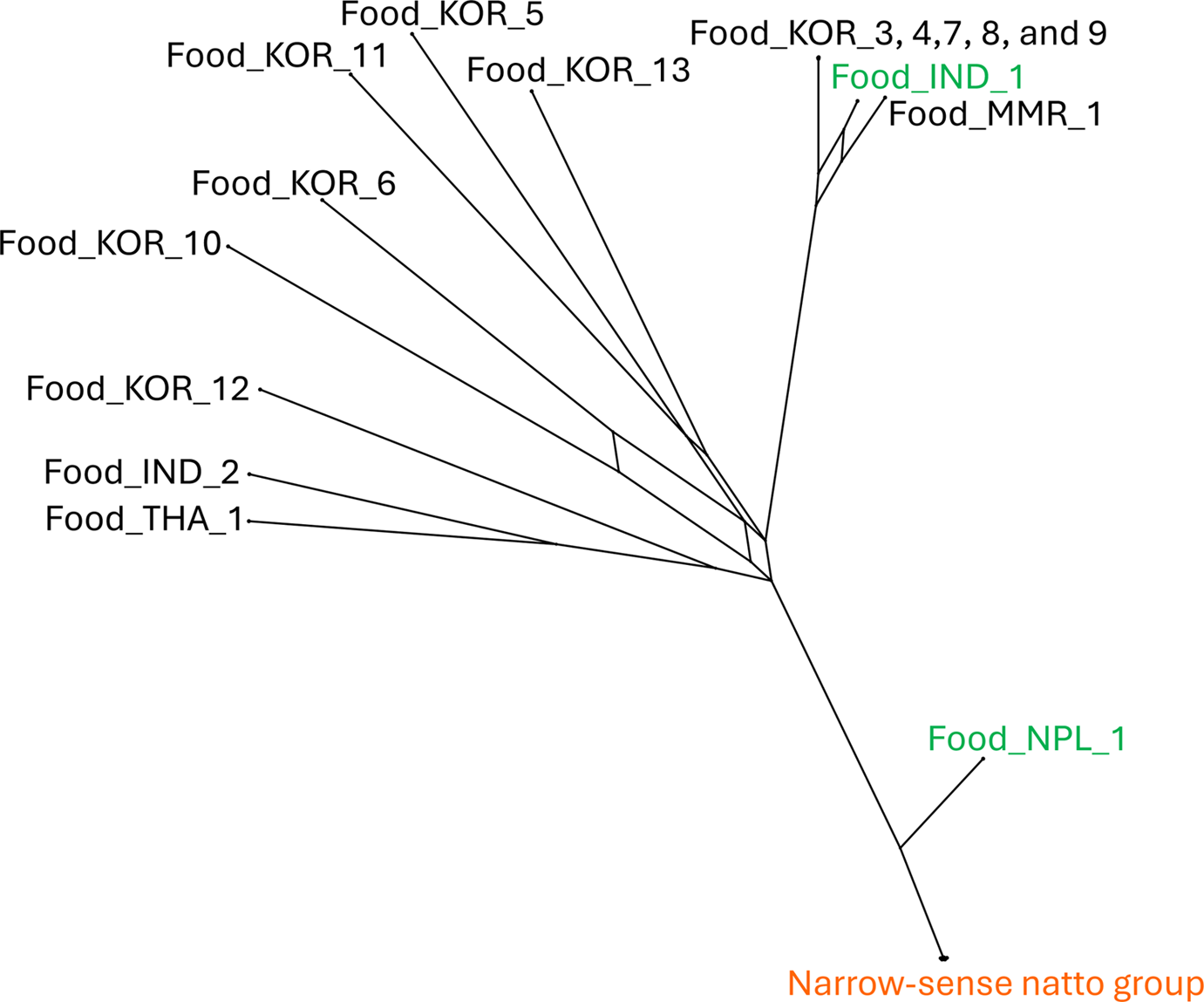
Phylogenetic network of 42 *B. subtilis* strains. The network was constructed from the core genome alignment using the Neighbor-Net algorithm. The tree-like portions of the network depict vertical, clonal inheritance, while the reticulate (box-like) structures indicate conflicting phylogenetic signals suggestive of recombination or horizontal gene transfer events.

### Identification and Functional Analysis of Group-Specific Accessory Genes

Based on the results above, we defined two additional operational groups for the subsequent functional comparison. The 15 strains phylogenetically related to the natto group but excluding the bekang strain are referred to as the “natto-related group”. The functionally defined cluster, consisting of the “narrow-sense natto group” (n=26) plus the single bekang isolate (Food_IND_1), is termed the “broad-sense natto group” (n=27 total).

To characterize the functional differences, we extracted genes specific to the “broad-sense natto group” (n=27) and the “natto-related group” (n=15). We defined “nearly group-specific genes” using a criterion of ≥80% presence within the target group and ≤20% presence in the comparison group. Using this criterion, we identified 184 nearly specific genes for the broad-sense natto group (Dataset 1) and 83 for the natto-related group (Dataset 2).

Gene Ontology (GO) and InterPro analyses of the broad-sense natto group-specific set (Figures 5, 6, Supplementary Figure 8) indicated an enrichment of DNA-binding proteins, enzymes involved in DNA rearrangement, various metabolic enzymes (hydrolases, transferases), and membrane-associated transporters (Dataset 3, Table 2). The analysis then focused on the natto-related group-specific gene set. The results of the GO analysis for the Biological Process (BP), Molecular Function (MF), and Cellular Component (CC) categories for this set are shown in Supplementary Figures 9, 10, and 11, respectively, with detailed annotations provided in Dataset 4. This analysis revealed that the natto-related group possesses its own distinct set of DNA-binding proteins, various metabolic enzymes with hydrolase or transferase activity, and membrane-localized transport enzymes. A selection of characteristic genes from this set is provided in Table 3.

**Figure 5.**
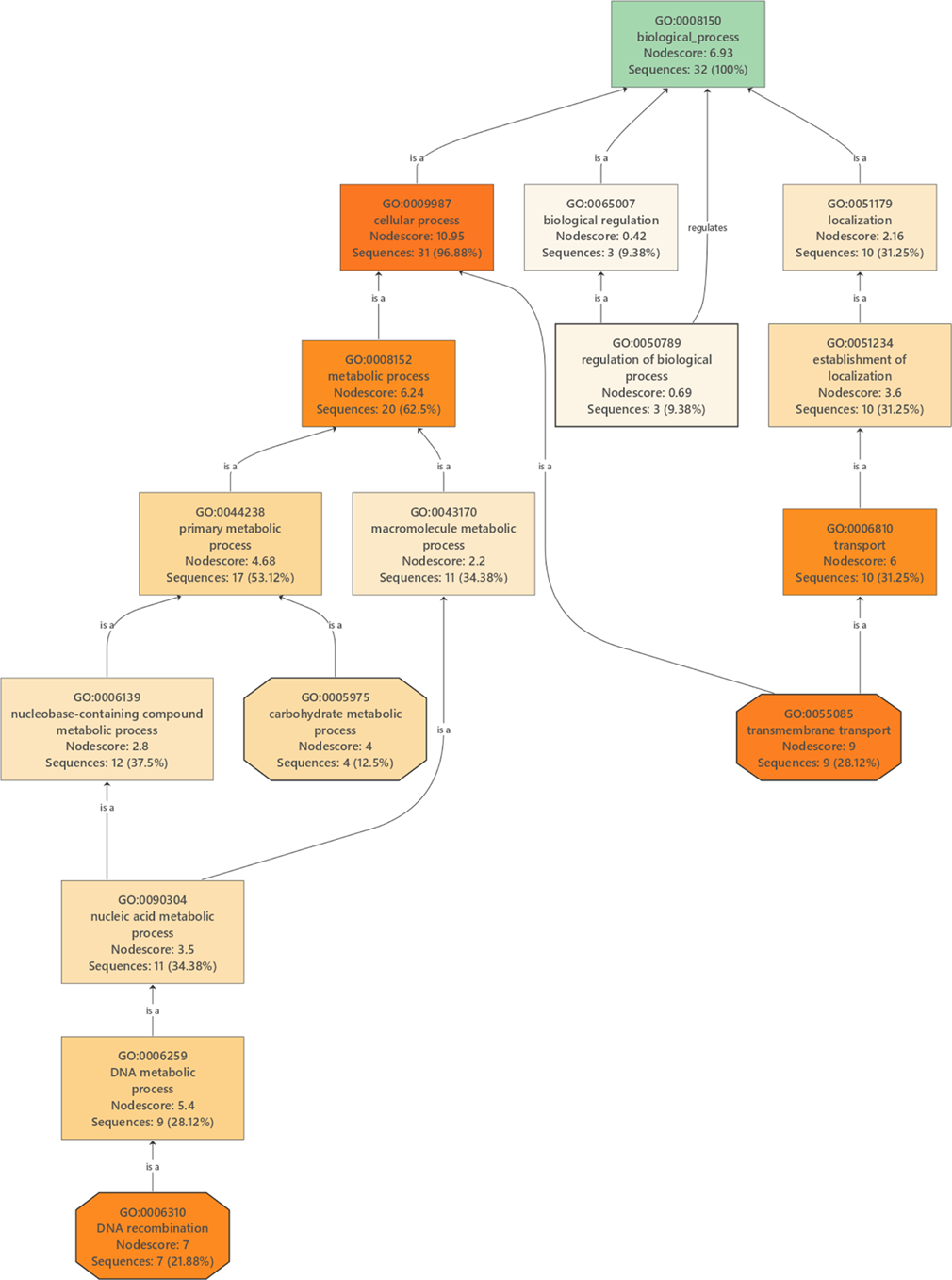
Gene Ontology (GO) enrichment analysis f for the “broad-sense natto group”-specific gene set. The figure shows a Directed Acyclic Graph of enriched GO terms, generated using OmicsBox. Each node (box) represents a GO term and displays its ID, description, NodeScore, and the number of associated gene sequences. The color intensity of each node is proportional to its enrichment significance, with more significantly enriched terms shown in shades of orange. The connecting lines represent hierarchical relationships between the terms.

**Figure 6.**
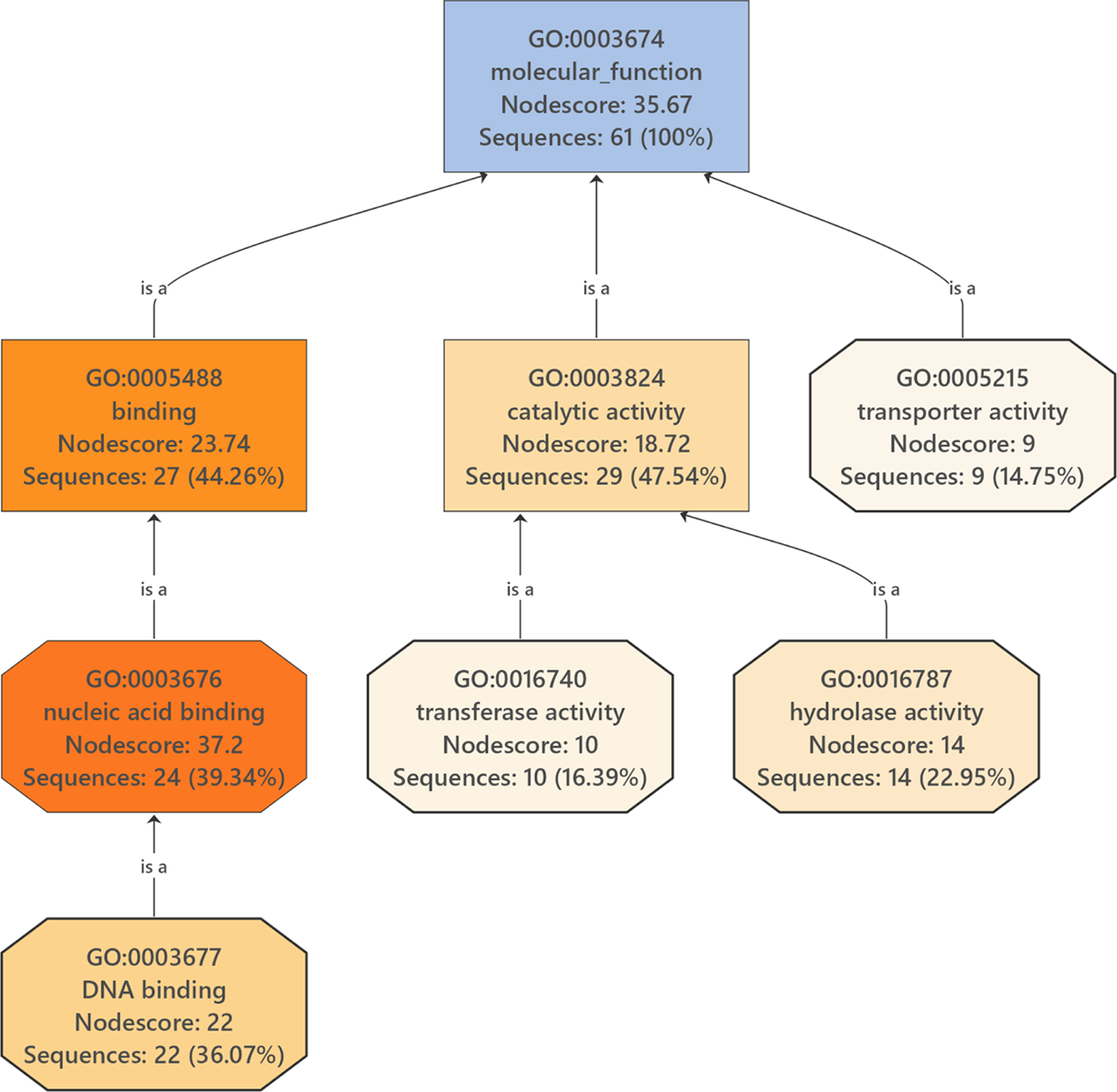
Gene Ontology (GO) enrichment analysis for the “broad-sense natto group”-specific gene set. The figure shows a Directed Acyclic Graph of enriched GO terms. Each box represents a GO term, displaying its ID, description, and the number of associated gene sequences. The color intensity of the boxes is proportional to the significance of enrichment, with more significant terms shown in shades of orange. The connecting lines represent hierarchical relationships between the terms.

**Table 2.**
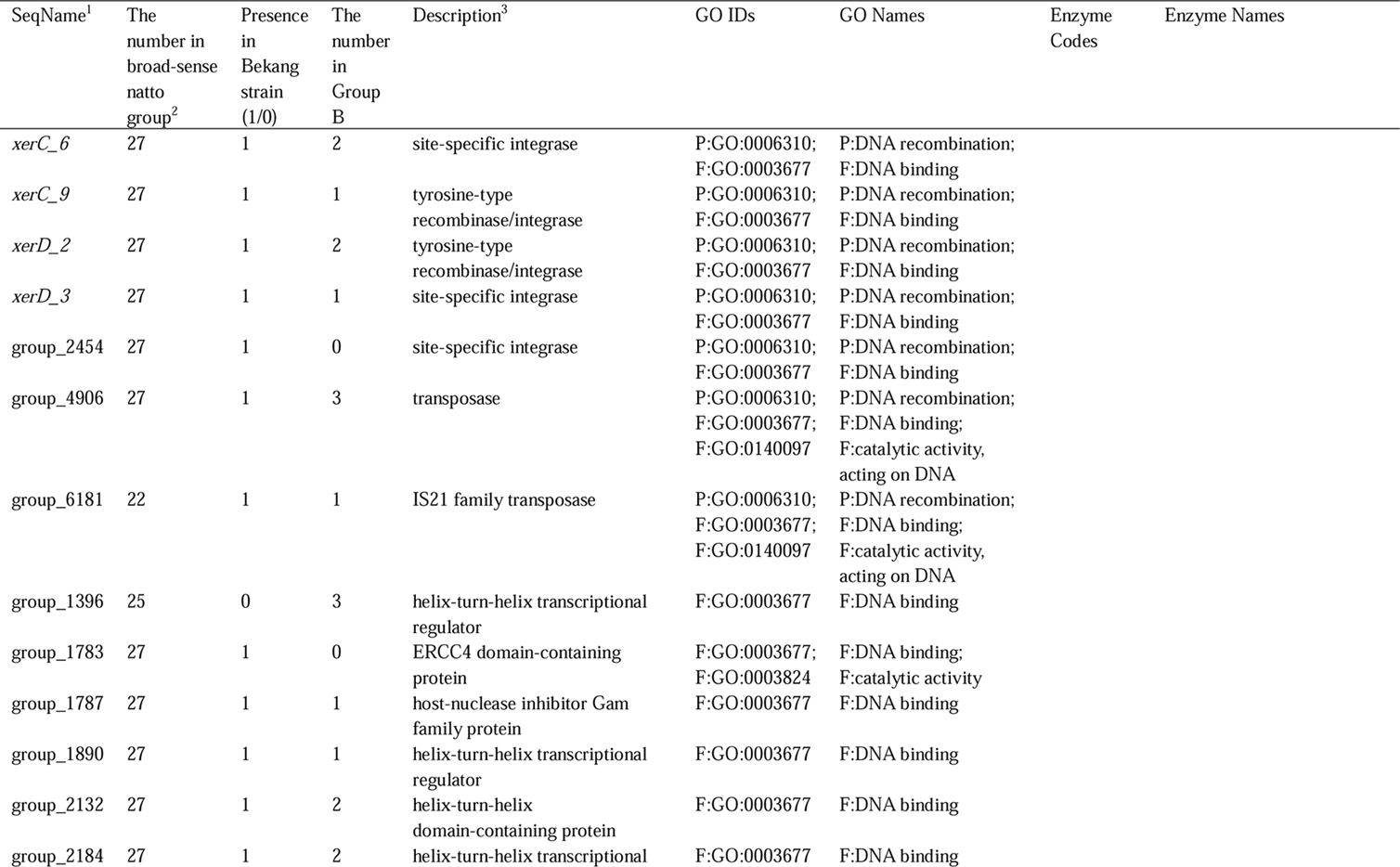
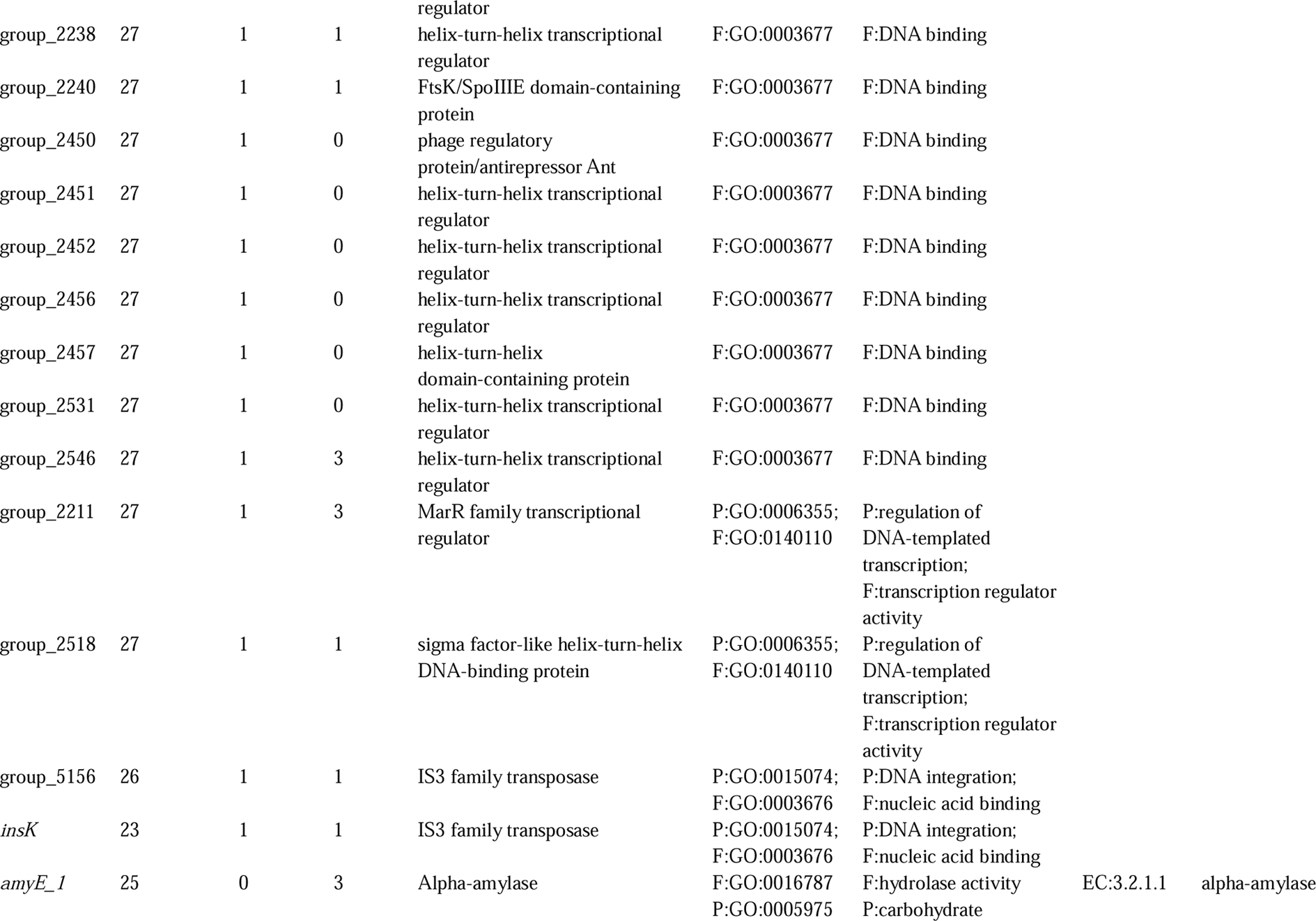
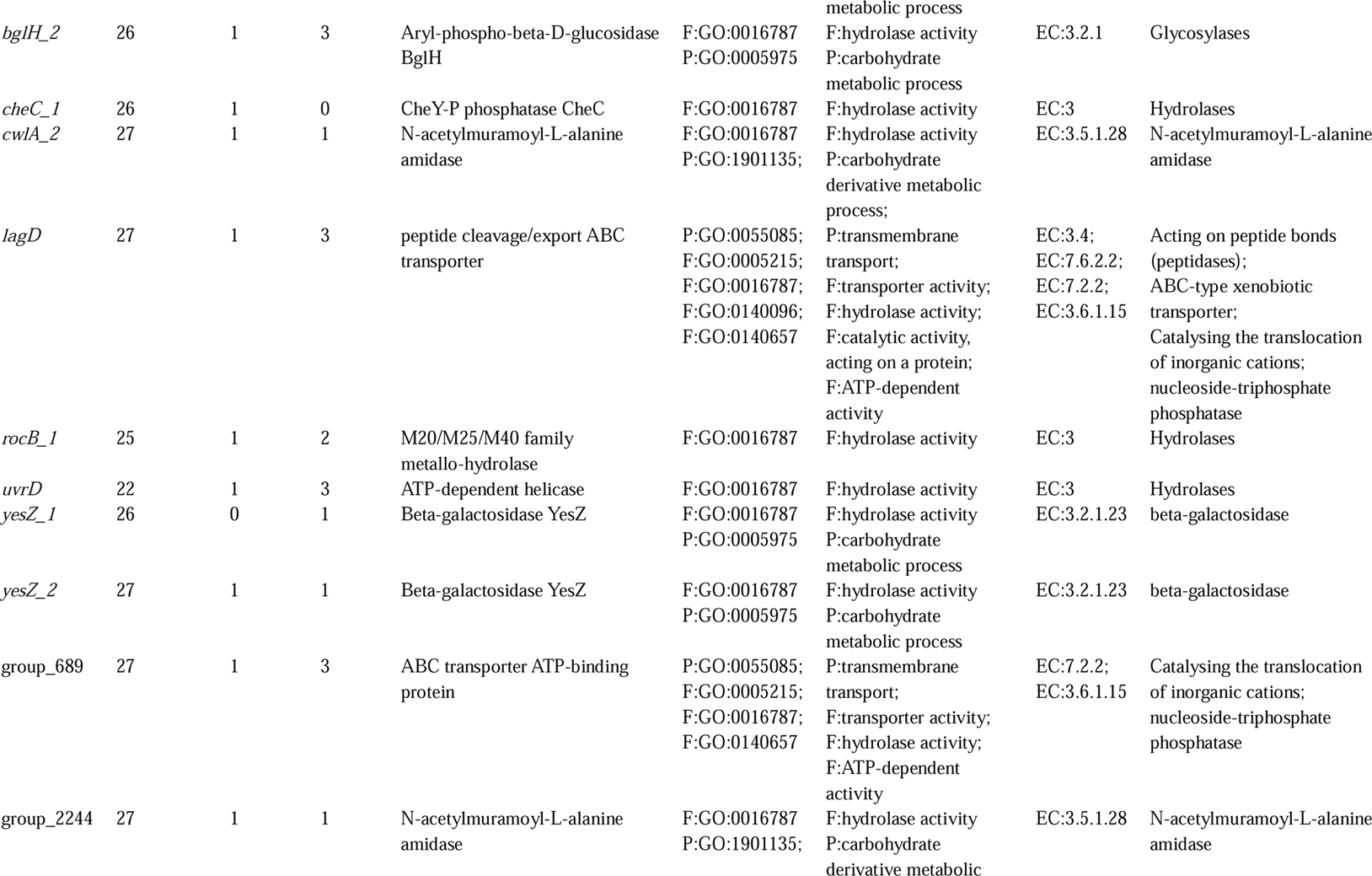
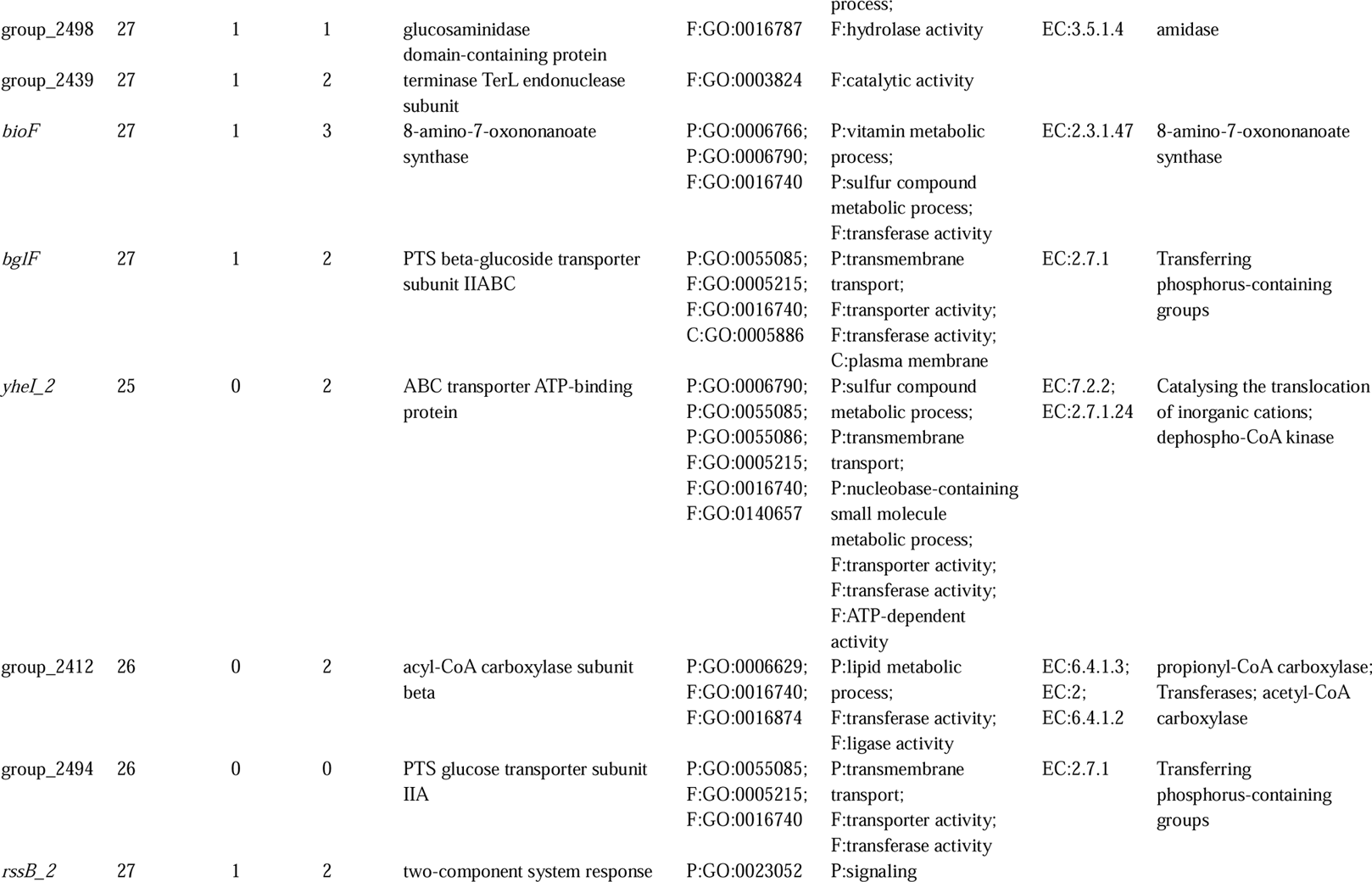
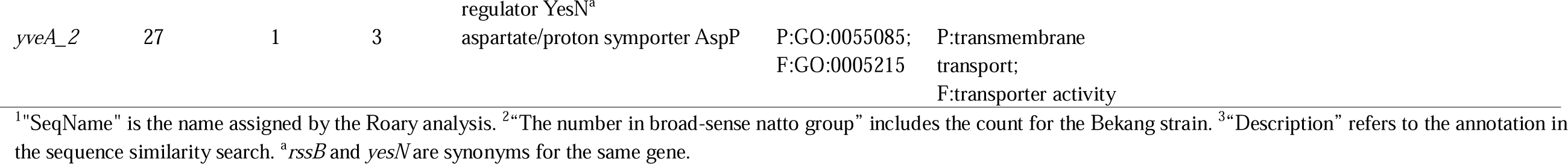
A selection of nearly “broad-sense natto group”-specific genes.

**Table 3.**
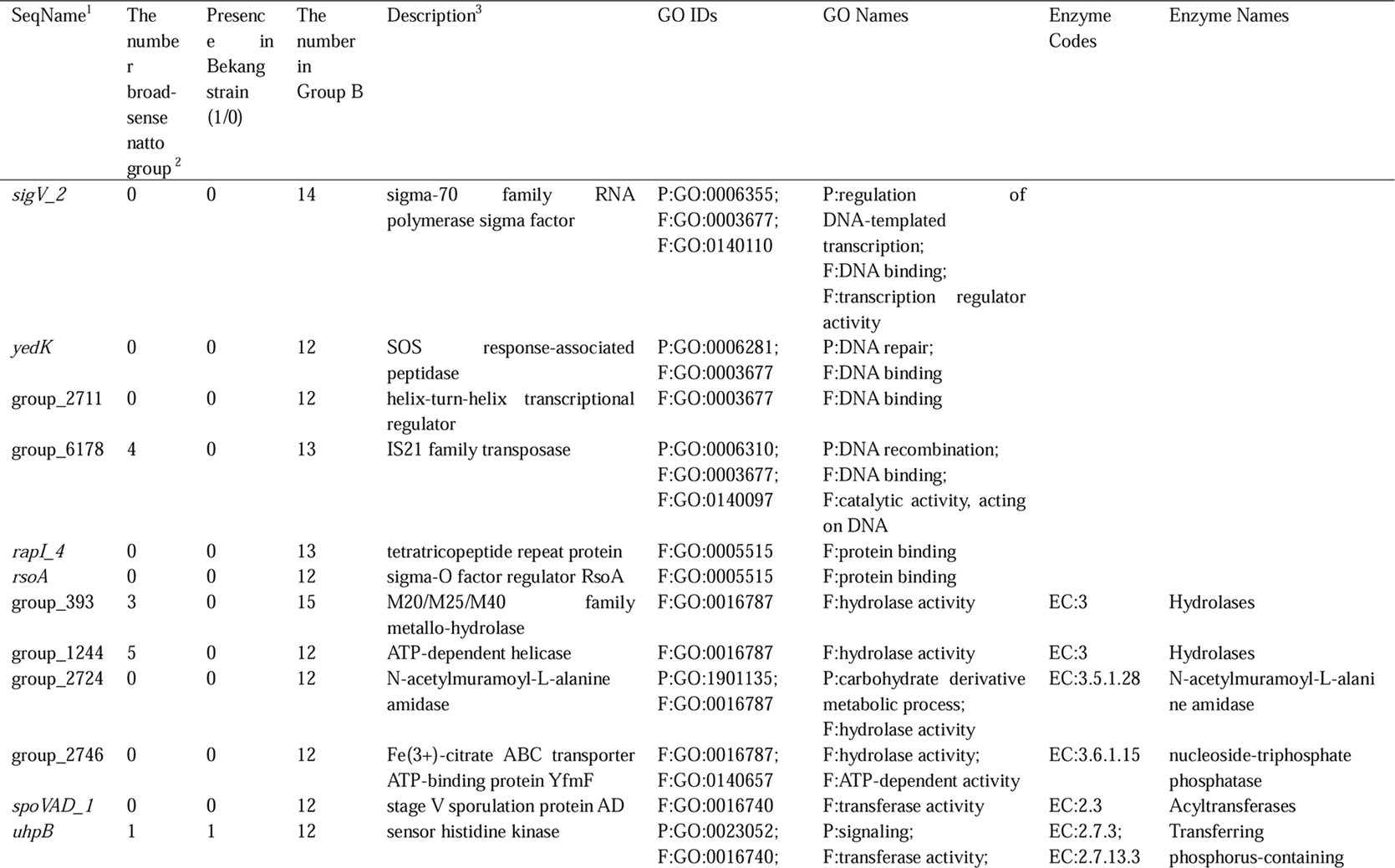
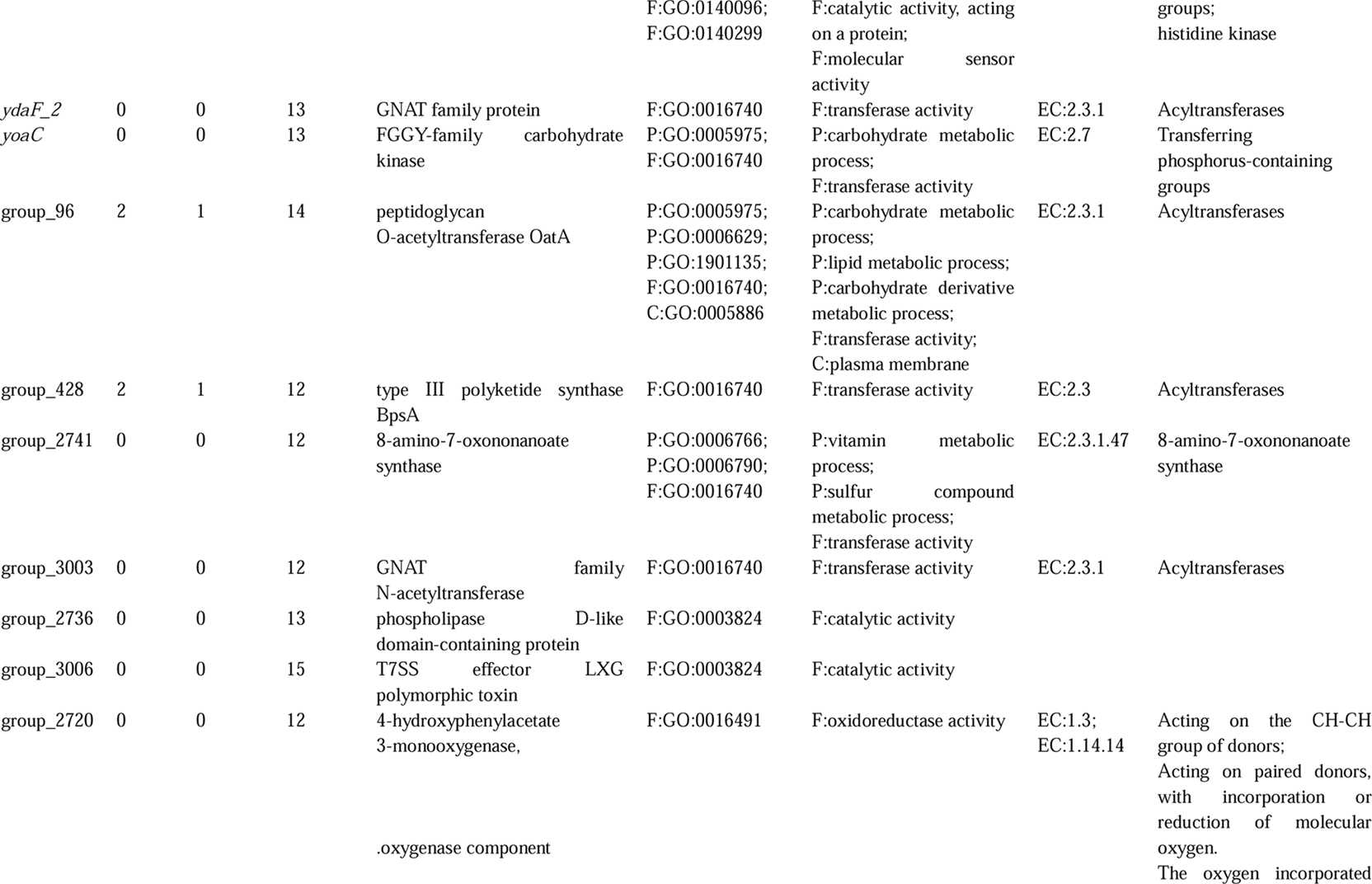
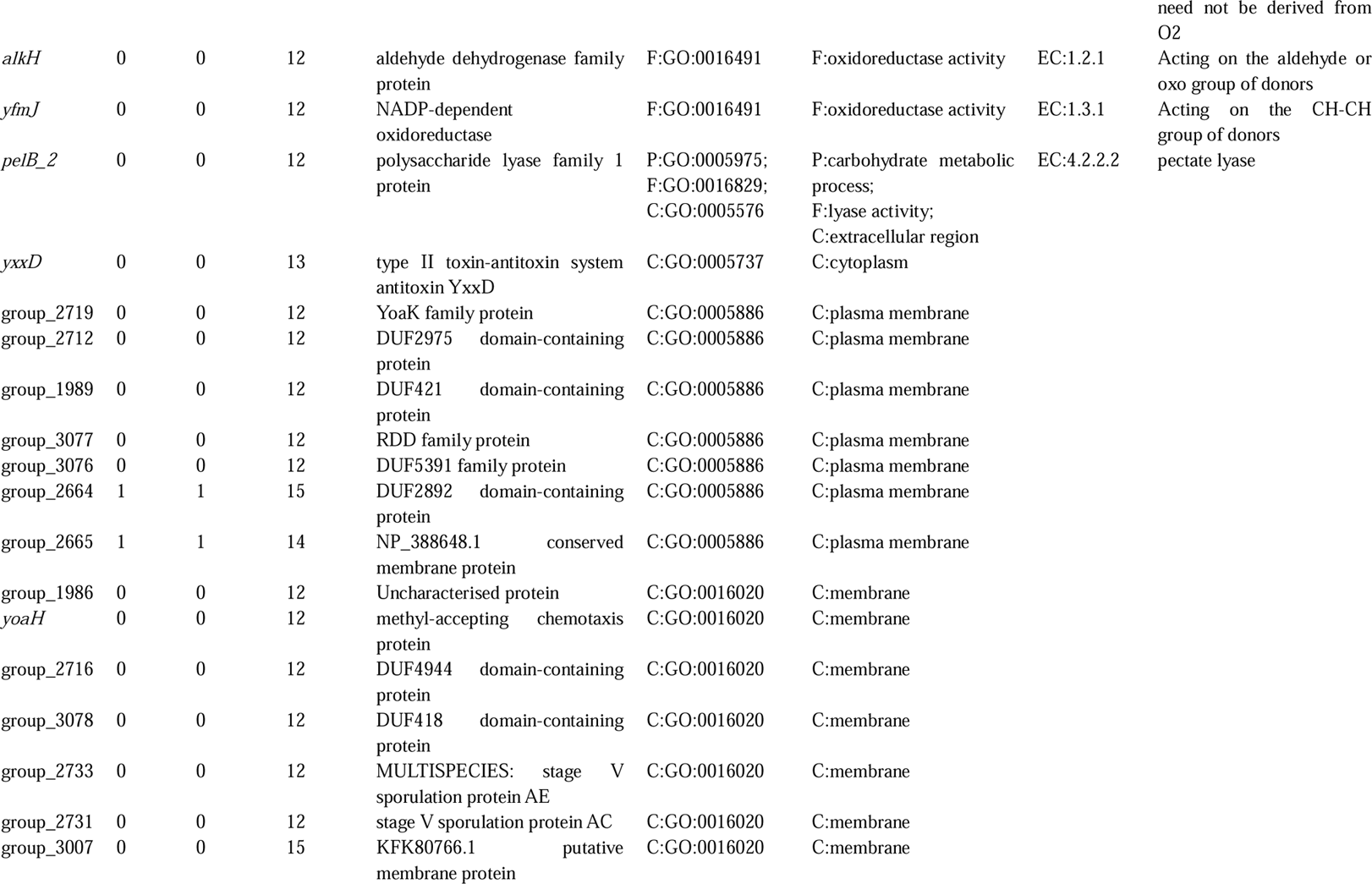
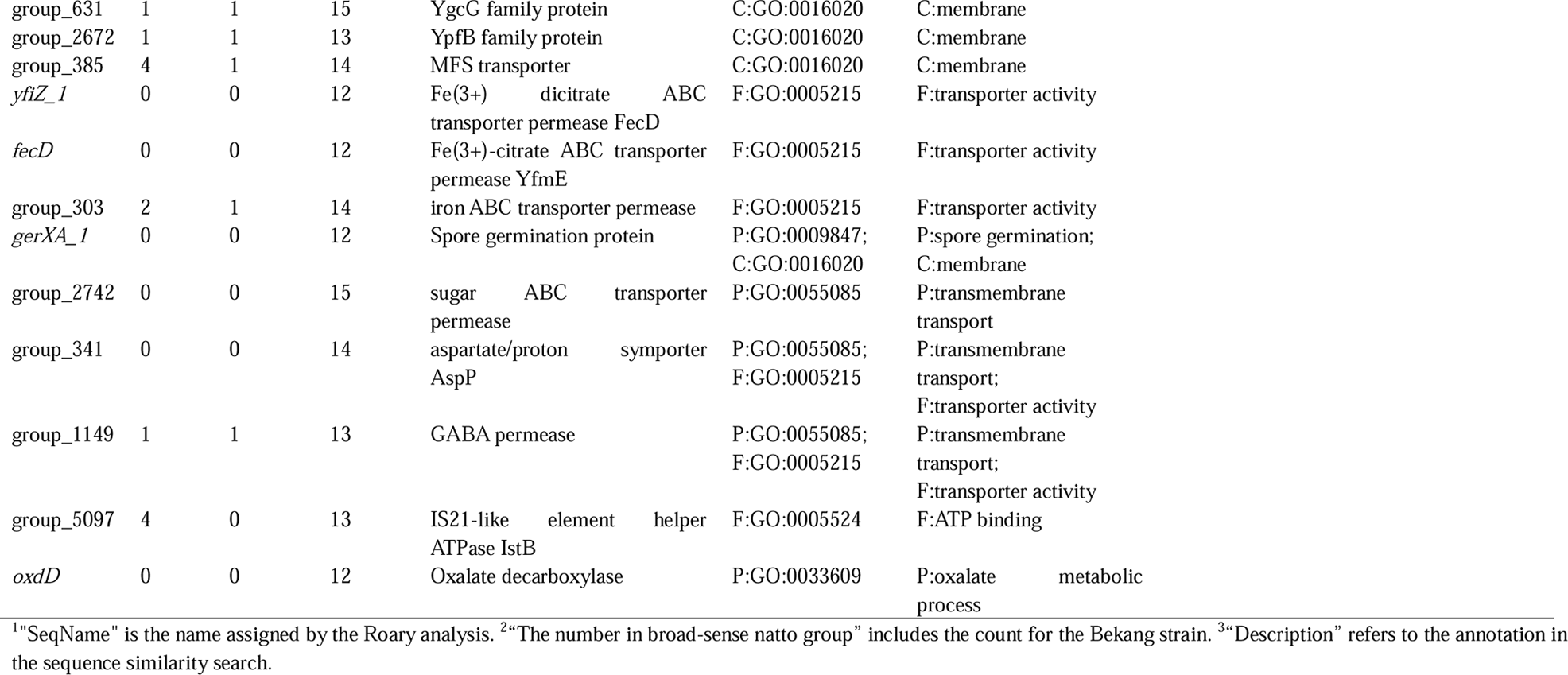
A selection of nearly “natto-related group”-specific genes.

We performed a simple sensitivity analysis to assess the impact of the chosen threshold for defining nearly group-specific genes. Altering the threshold (to 70%/30% or 90%/10%) predictably changed the total number of genes identified (Datasets 5, 6, 7, 8, 9, 10, 11, and 12). However, the major functional categories enriched in the resulting gene sets for both the broad-sense natto group and the natto-related group remained largely consistent, confirming the robustness of our functional interpretation based on the 80%/20% threshold.

Analysis of particularly interesting genes revealed the following: “broad-sense natto group” strains typically possess two highly similar copies of *rocB* (arginine degradation)^34^ and *yveA* (aspartate-proton symporter)^35,36^, whereas the “natto-related group” strains contain only one. This group also possesses a duplicate copy of *yesN*, a two-component system response regulator^37^. The *amyE* gene (alpha-amylase)^38^ also showed duplication in most “narrow-sense natto group” strains, though the bekang isolate (Food_IND_1) contained only a single copy. The *bglF* homolog (transport of β-glucosides)^39^ in the “broad-sense natto group” is significantly longer (609 amino acids) than the version in the “natto-related group” (461 amino acids). In contrast, the *bglH* (degradation of β-glucosides)^39^ and *bioF* (biotin synthesis)^18,40^ genes in the “broad-sense natto group” are notably shorter, truncated versions compared to the full-length homologs found in the “natto-related group”. Among the results for the natto-related group-specific gene set, two genes are of particular interest: *pelB* (Pectate lyase)^41^ and *sigV* (sigma factor V)^42^. For *pelB*, no highly similar sequences were detected in the broad-sense natto group. Furthermore, the *pelB* gene exhibited remarkable diversity even within the natto-related group itself. Only the homolog from the kinema-producing *B. subtilis* strain (Food_IND_2) appeared to be full-length, encoding a protein of 304 amino acids. In contrast, the versions found in other natto-related group strains were considerably shorter, ranging from 166 to 170 amino acids, suggesting that they are either truncated or functionally distinct. The other gene of interest, *sigV*, exhibited copy number variation, with natto-related group strains typically possessing two copies while broad-sense natto group strains have only one.

The loss of function in the flagellar gene *fliF,* previously noted as a characteristic of *B. subtilis* var. *natto*^18^, was observed in “narrow-sense natto group” strains but not in the bekang isolate, indicating it is not a universal trait of the “broad-sense natto group”. Finally, the gene presence/absence analysis revealed no differences between the “broad-sense natto group” and the “natto-related group” for the key genes associated with polyglutamic acid (PGA) synthesis (*pgsBCA*) or its main regulators^3^. Furthermore, the synteny of the genomic region containing these PGA-related genes was found to be highly conserved across all analyzed strains. This finding is consistent with previous comparative study^18^, which also failed to identify clear differences within the PGA synthesis operon itself that could explain the phenotypic variations in PGA production between natto and related strains.

## Discussion

In this study, we present a compelling case study of a single *B. subtilis* strain from Indian bekang (Food_IND_1) that exhibits a striking accessory gene similarity to the phylogenetically distant Japanese *B. subtilis* var. *natto* clade (“narrow-sense natto group”). This core finding highlights a discrepancy between systematic relatedness (based on core genes) and functional similarity (based on accessory genes) at the boundary of the natto clade. Our quantitative analysis using Jaccard distance clearly positioned the bekang strain as the functional nearest neighbor to the narrow-sense natto group, while the systematic nearest neighbor from Nepal (Food_NPL_1) was functionally more distant (Figure 2). This finding is particularly intriguing given that a defining characteristic of Japanese natto is its intense PGA-derived viscosity—a feature that distinguishes it from most other *B. subtilis*-fermented soybean products. As noted in the Introduction, bekang is also reported to be a highly viscous and stringy product^7^. Bekang is a traditional food consumed not only in Mizoram, India, but also in the adjacent Chin State of Myanmar. The shared genomic profile leads to the hypothesis that this specific bekang-producing lineage may also possess a notable capacity for PGA production. Indeed, a previous study reported substantial PGA production in other bekang-isolated strains, supporting this hypothesis^7^. Although the strains analyzed in that study were different from Food_IND_1, the potential for high PGA production presents a compelling candidate for the key functional trait linked to the conserved accessory gene repertoire (“broad-sense natto group” signature) we observed.

How did the Food_IND_1 strain come to share an adaptive module with distant natto strains? We propose two competing, and equally plausible, evolutionary scenarios. The first hypothesis is horizontal gene transfer (HGT)^43^. We propose that a “natto-type” adaptive module—a large, co-evolved set of genes conferring advantageous traits for soybean fermentation—was transferred as a “plug-and-play” system, allowing for rapid adaptation to the soybean niche^44^. This is plausible, as *B. subtilis* is known for its natural competence^45^. Our network analysis, which detected genetic exchange between the bekang strain and a Myanmar strain, suggests this lineage may be part of a broader network of gene flow. The alternative hypothesis involves selective retention of ancestral traits and differential gene loss^46,47^. This scenario posits that a common ancestor of both the “narrow-sense natto group” and the “natto-related group” possessed this “soybean-adaptation” gene set. Subsequently, this module was lost in most “natto-related” lineages (like the Nepalese strain) but was selectively retained in both the bekang and natto lineages due to similar selective pressures encountered in viscous soybean fermentation niches. At present, our data do not allow us to definitively distinguish between these two scenarios. Both remain viable explanations for the observed genomic paradox.

Our analysis also highlighted the remarkable genetic homogeneity among the Japanese *B. subtilis* var. *natto* strains (the “narrow-sense natto group”). This is likely attributable to the modern industrial practice of using a limited number of elite starter cultures (spores) selected for high PGA productivity^4^, a practice that imposes low evolutionary pressure during fermentation and creates an artificial bottleneck. However, the presence of phylogenetically similar strains within this group isolated from natural environments (e.g., the deep-sea isolate Env_JPN_1^12^) suggests the lineage is not purely an industrial artifact. This might be explained by the “escape” of resilient industrial spores back into diverse ecosystems, or potentially that the lineage was already naturally homogeneous before widespread industrialization.

The distribution of food-derived (Food) and environmental (Env) strains within our focused 42-strain dataset presents a complex picture that resists simple interpretation regarding selective pressures. The “natto-related group” (n=16) consists entirely of food isolates, reflecting the study’s focus on strains from fermented foods that are phylogenetically close to the natto clade. Conversely, the highly homogeneous “narrow-sense natto group” (n=26) itself contains strains isolated from environmental sources (e.g., Env_JPN_1). The origin and history of these environmental strains clustering within the core natto clade are unknown; they could represent strains escaped from industrial settings, strains from natural populations closely related to those used industrially, or other scenarios. Given these significant uncertainties, it is difficult to definitively conclude how food fermentation history versus natural environmental pressures have shaped the observed genomic patterns based solely on the Food/Env labels. What remains clear, however, is the existence of the conserved “natto-type” accessory gene module. Its presence in both the core natto clade (with its mix of Food and Env isolates) and the phylogenetically distant, food-derived bekang isolate suggests this adaptive toolkit is not merely a product of modern industrial selection but represents a pre-existing, successful strategy for utilizing soybean substrates.

These findings strongly support a model of polygenic adaptation, in which characteristic traits such as efficient soybean substrate utilization are not dictated by a single gene but by a large, conserved set of accessory genes acting in concert. The “broad-sense natto group”-specific gene set represents one such successful polygenic module.

Finally, this study has several primary limitations. First, and most critically, our central finding regarding the natto-bekang connection is based on the genomic analysis of a single bekang isolate (n=1). Future studies involving the genomic analysis of a broader collection of *B. subtilis* strains from bekang are crucial to determine whether the conserved accessory repertoire we observed in Food_IND_1 is a common feature. Furthermore, this investigation should be extended to the diverse *B. subtilis* strains from Myanmar’s fermented soybean foods. The Bamar people of Myanmar represent another major population—alongside the Japanese and Koreans—with a rich tradition of producing *B. subtilis*-fermented soybean foods, making the region a critical and largely unexplored nexus for this bacterium’s diversity. Our discovery of genetic exchange between the Indian bekang and Myanmar pe poke strains further underscores the importance of this region. Such expanded research will be essential to map the complex pathways of gene flow and understand the adaptive radiation of these bacteria across Asia. Second, our study is based on *in silico* genomic data. We have identified a set of candidate genes, but we have not shown that they are actively expressed. Future work, such as transcriptomics, is essential to confirm whether these genes are truly being expressed during soybean fermentation and contribute to the adaptive phenotype. A further limitation is that many genomes used were draft assemblies. This makes the reliable analysis of repetitive sequences like IS elements, which are known to be abundant in natto strains and act as important evolutionary drivers, technically challenging. This remains an important avenue for future research using complete genomes.

## Methods

### Genome Data Acquisition and Quality Control

The 55 *B. subtilis* strains used in this study are detailed in Table 1. This initial set was selected to explore the diversity surrounding *B. subtilis* var. *natto*, including strains identified through sequence similarity searches using *B. subtilis* var. *natto* sequences as queries and from relevant literature. The key bekang isolate (Food_IND_1, Miz-8) was not a culture collection strain, but was reported as being directly isolated from traditional bekang in Mizoram, India^6^. For genomes where assemblies were not publicly available, raw reads were retrieved from public read archives and assembled de novo using SPAdes (version 4.2.0)^48^. Assembled contigs shorter than 201 bp were subsequently removed using the seq -m 201 option in SeqKit (version 2.8.2)^49^. The quality of all genome sequences was assessed by calculating completeness and contamination values using CheckM2 (version 1.1.0)^50^. To ensure consistent annotation across all datasets, all genome sequences were re-annotated using Prokka (version 1.14.6)^51^. The resulting GFF, GenBank, and FAA files are available from https://drive.google.com/file/d/1dRA0wjLvmyoJHyVJF8F1l4Pl5rchX7GS/view.

### Pangenome and Phylogenetic Analysis

Using the GFF files generated by Prokka as input, a pangenome analysis was conducted with Roary (version 3.13.0)^31^ using the options -e -n -v -r. The resulting gene presence/absence matrices for both the full set of 55 samples (gene_presence_absence_55_samples.csv) and the subset of 42 samples (gene_presence_absence_42_samples.csv) can be accessed at https://drive.google.com/file/d/1dRA0wjLvmyoJHyVJF8F1l4Pl5rchX7GS/view. Core-gene phylogenetic trees were constructed from the core_gene_alignment_55_samples.aln and core_gene_alignment_42_samples.aln files (available from https://drive.google.com/file/d/1dRA0wjLvmyoJHyVJF8F1l4Pl5rchX7GS/view) generated by Roary. For building a maximum-likelihood phylogenetic tree, IQ-TREE 2 (version 2.2.2.3)^52^ was used with the options -m MFP -bb 1000 -alrt 1000. The -m MFP option enables automatic substitution model selection. The best-fit substitution model, as determined by ModelFinder under the Bayesian Information Criterion (BIC), was GTR+F+I+R5 for the 55-strain analysis and GTR+F+I+R2 for the 42-strain analysis. The resulting phylogenetic tree, including support values, was visualized using FigTree (version 1.4.4) (http://tree.bio.ed.ac.uk/software/figtree/). To generate a figure that merges the phylogenetic tree with the gene presence/absence matrix (Supplementary Figure 2), we used a modified version of the roary_plots.py script, which is provided at https://github.com/sanger-pathogens/Roary.git. This modified script, roary_plots_2.py (Dataset 13), takes the treefile from IQ-TREE 2 and the gene_presence_absence.csv file from Roary as input. The primary modification to this script was the implementation of midpoint rooting for the phylogenetic tree.

### Cluster Analysis Based on Gene Presence/Absence Profiles

To perform cluster analysis based on functional profiles, the gene_presence_absence.csv files generated by Roary (gene_presence_absence_55_samples.csv for the 55-strain analysis and gene_presence_absence_42_samples.csv for the 42-strain analysis) were directly analyzed using a custom Python script, one_step_cluster.py (Dataset 13). This script calculates pairwise distance matrices using several metrics, including Jaccard, Sørensen-Dice, Hamming, and Euclidean distances. For this study, we selected the Jaccard distance metric, as it appropriately ignores joint absences (“double zeros”) in binary presence/absence data. Based on the calculated Jaccard distance matrices for both the 55 strains and the focused 42 strains, the one_step_cluster.py script performed hierarchical clustering and generated the resulting heatmaps and dendrograms. This analysis was employed as a quantitative and exploratory map to assess functional similarity, rather than to define statistically discrete groups.

### Visualization of Synteny Blocks

To visualize synteny blocks among the three selected chromosome-level genomes, Easyfig (version 2.25)^53^ was used. This analysis required three GenBank annotation files and two BLAST^54^ comparison files as input. The annotation files were the GenBank-formatted outputs generated by Prokka. The two required BLAST files were generated from sequential pairwise blastn searches: the first search compared genome 1 (query) against genome 2 (database), and the second compared genome 2 (query) against genome 3 (database). For both searches, the output was specified in tabular format using the -outfmt 6 option. Using these three GenBank files and two BLAST files as input, Easyfig was run with its default parameters to generate the comparative visualization.

### Phylogenetic Network Analysis

A phylogenetic network was constructed from the core gene alignment to investigate potential reticulate evolutionary events, such as recombination. The analysis was performed using SplitsTree (version 6.4.16)^55^ with default settings. The input file was the core_gene_alignment_42_samples.aln or core_gene_alignment_55_samples.aln generated by Roary, which was renamed with a .fasta file extension prior to the analysis.

### Identification and Extraction of Group-Specific Accessory Genes

For the identification of “broad-sense natto group”-specific and “natto-related group”-specific genes, “nearly group-specific genes” were first defined using a criterion of ≥80% presence within the target group and ≤20% presence in the comparison group. Alternative thresholds, such as 70%/30% and 90%/10%, were also evaluated to assess the robustness of the findings. The analysis was performed on the gene_presence_absence_42_samples.csv file from the 42-strain Roary analysis (comprising the “narrow-sense natto group”, the bekang strain, and the “natto-related group”). Custom scripts, analyze_genes_A_specific.py and analyze_genes_B_specific.py (Dataset 13), were executed to parse this file and extract the names of all genes that met the defined criteria for the “broad-sense natto group” and “natto-related group”, respectively. The resulting tabular output was then processed with another custom script, process_csv.py (Dataset 13), to create a list containing only the primary gene names from the leftmost column. Finally, a third script, process_fasta.py (Dataset 13), utilized this list of gene names to retrieve the corresponding amino acid sequences in FASTA format from the multiple per-sample protein files (.faa) that were generated by Roary and stored in a specified directory.

### Functional Annotation

Functional annotation of the “broad-sense natto group”-specific and “natto-related group”-specific protein sets (.faa files) was performed using OmicsBox (version 3.4.6; GO version March 16, 2025). The workflow consisted of several integrated steps. First, a similarity search was conducted against the Diamond BLAST database (2024-11-20 release) using the blastp mode^56^. This search was taxonomically filtered to *Bacillus* (taxid: 1386) with an E-value cutoff of 1.0E-3. Next, protein domains and families were identified by running InterProScan (version 5.72-103.0)^57^. Finally, GO terms were mapped and assigned to each sequence within OmicsBox based on the combined results of the previous steps^58,59,60^. For the final annotation, results were filtered with an E-value-Hit Filter of 1.0E-6 and a Hit Filter of 500.

### Use of Generative AI

The generative AI models Gemini and ChatGPT were used to improve the language and clarity of the manuscript and to assist in writing the Python scripts.

## Acknowledgements

We are especially grateful to Dr. Myat Htoo San of the National Institute of Advanced Industrial Science and Technology (AIST), Hokkaido, Japan, and her friend, Ms. Zing Cin Tial, a member of the Chin community from Hakha, Chin State, Myanmar. They kindly provided valuable information and cultural insights into the traditional production of bekang in Chin State, Myanmar, for which there is little published literature. We also thank Drs. Taiki Futagami and Hisanori Tamaki of the Faculty of Agriculture, Kagoshima University, and Drs. Masatoshi Goto and Genta Kobayashi of the Faculty of Agriculture, Saga University, for guiding us into the research of fermented foods.

## Author Contributions

K.S. and Y.N. conceived and designed the study, performed the bioinformatic analyses, and interpreted the data. Y.N. wrote the original manuscript draft, and K.S. reviewed and edited the manuscript. Both authors read and approved the final manuscript.

## Data Availability Statement

The primary genome sequence data analyzed in this study were retrieved from publicly available databases; specific accession numbers for all strains are detailed in Table 1. The key data files generated during the computational analysis, including the gene presence/absence matrices, can be accessed at https://drive.google.com/file/d/1dRA0wjLvmyoJHyVJF8F1l4Pl5rchX7GS/view. All custom Python scripts developed for this work are available as Dataset 13.

