## Supplementary Information for "Conserved accessory genes link a phylogenetically distinct *Bacillus subtilis* strain from Indian bekang to the Japanese natto clade"

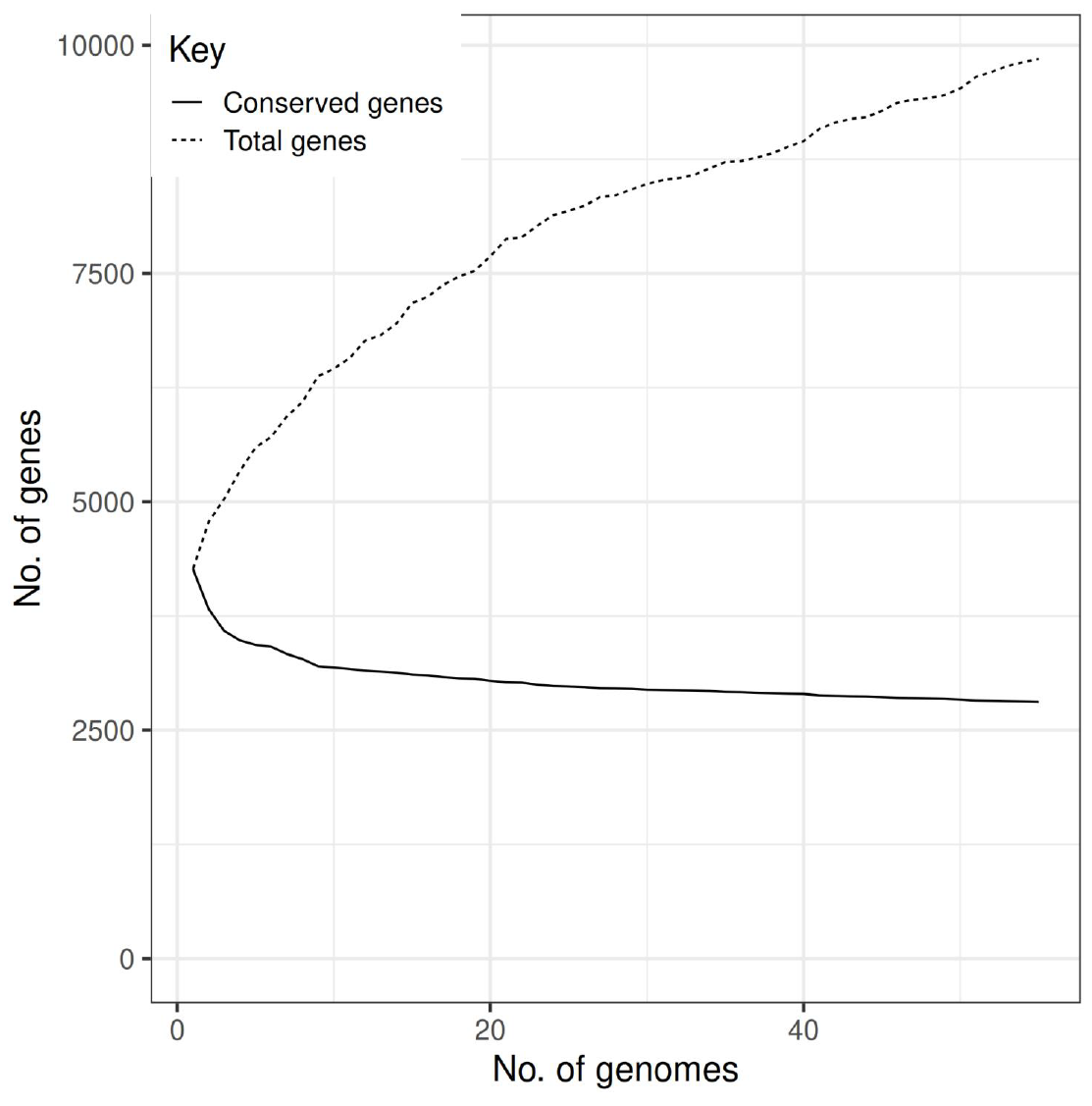
**Supplementary Figure 1.** Pangenome analysis of the 55 *Bacillus subtilis* strains. The plot shows the number of genes (y-axis) as a function of the number of genomes analyzed (x-axis). The upper curve represents the pangenome size (total genes), and the lower curve represents the core genome size (conserved genes).

**Supplementary Figure 2.** Core-genome-based phylogeny and accessory gene presence/absence matrix of the 55 *B. subtilis* strains. The phylogenetic tree on the left was constructed from a core genome alignment of 55 strains using the maximum likelihood method and is presented as midpoint-rooted. The matrix on the right, generated by Roary, visualizes the pangenome, which consists of 9,848 gene clusters. Each row corresponds to a strain in the tree, and each column represents a single gene cluster. Dark blue cells indicate the presence of a gene cluster in a strain, while white cells indicate its absence. The "narrow-sense natto group" (n=26) is indicated by an orange line. Key strains discussed in the text, the systematic nearest neighbor (Food_NPL_1) and the functional nearest neighbor (Food_IND_1), are highlighted with green squares on their labels.

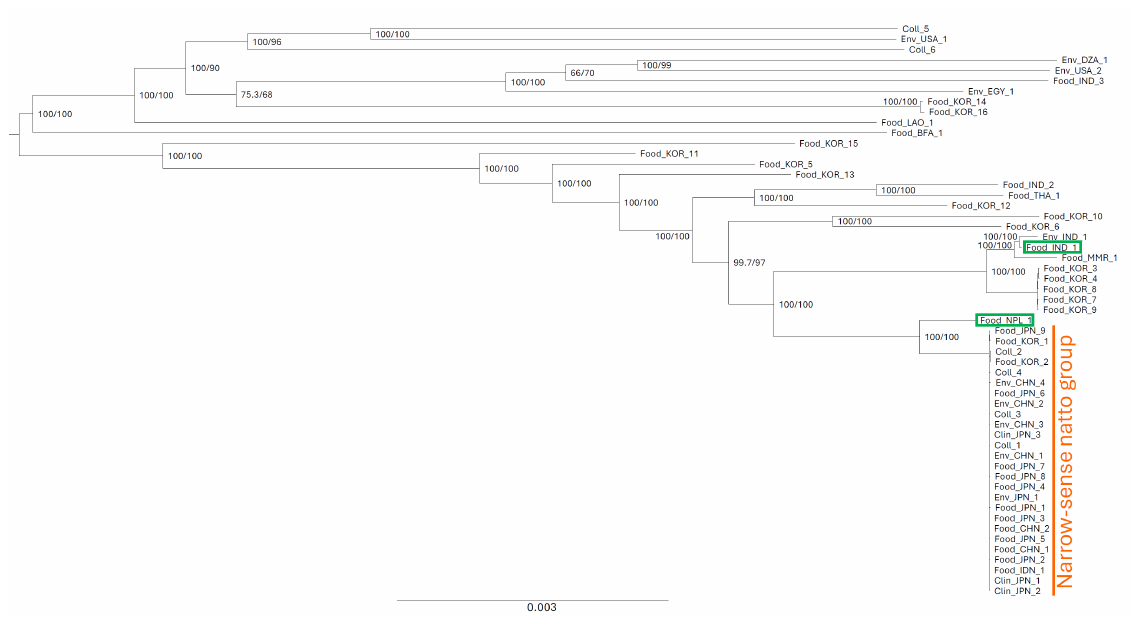

**Supplementary Figure 3.** Core-genome-based phylogenetic tree of the 55 *Bacillus subtilis* strains with support values. This phylogenetic tree was constructed from the core genome alignment of the 55 strains using the maximum-likelihood method with IQ-TREE 2. The tree is displayed using midpoint rooting. Numbers at the nodes indicate support values from SH-aLRT and UFBoot. Support values for tightly clustered strains are not shown for clarity. The "narrow-sense natto group" (n=26) is indicated by an orange line. Key strains discussed in the text, the systematic nearest neighbor (Food_NPL_1) and the functional nearest neighbor (Food_IND_1), are highlighted with green squares on their labels.

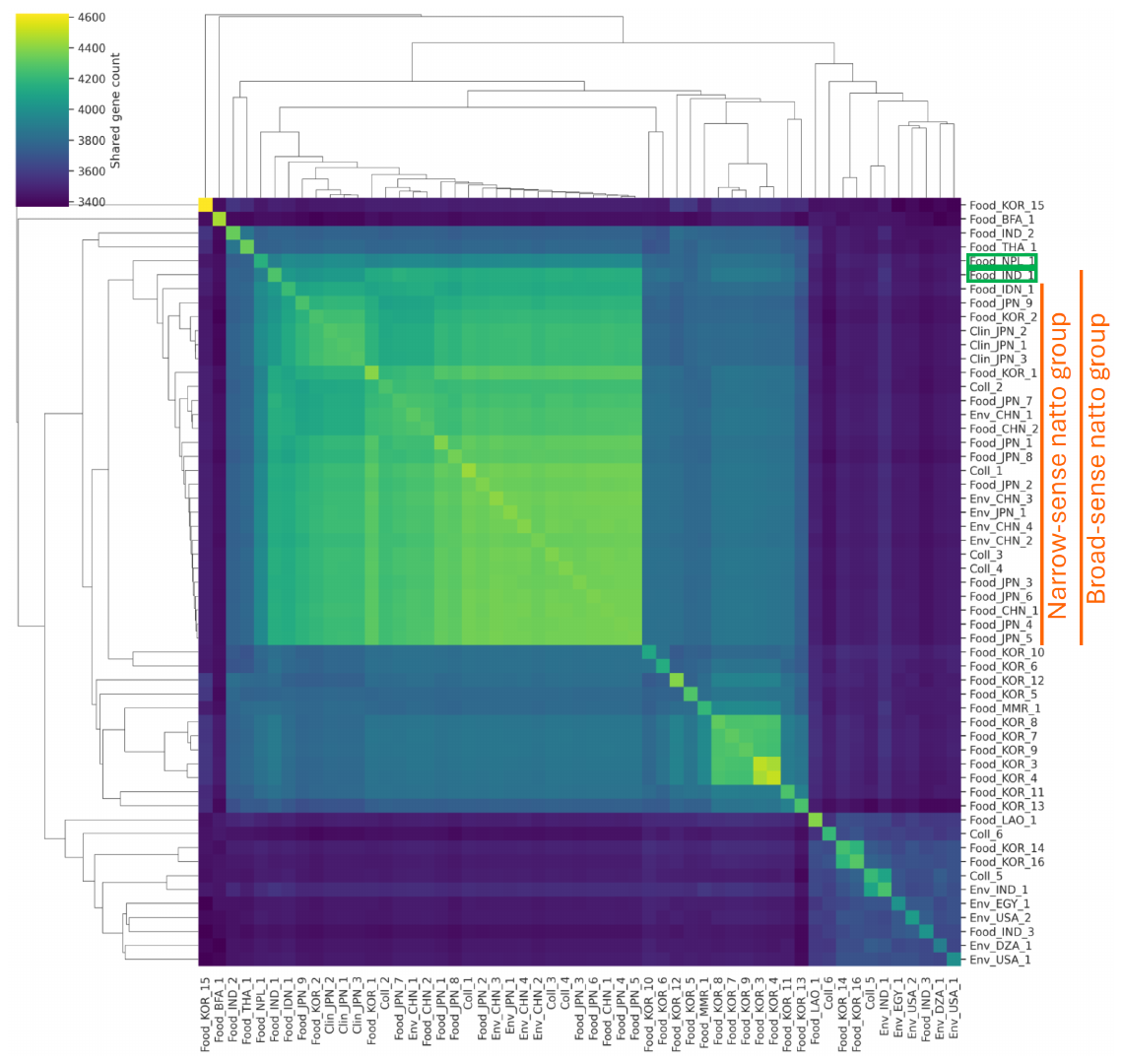

**Supplementary Figure 4.** Hierarchical clustering of 55 *B. subtilis* strains based on accessory gene profiles. The heatmap shows the pairwise shared gene counts between all 55 strains. Hierarchical clustering was performed using Jaccard distance calculated from the binary presence/absence matrix. The "narrow-sense natto group" (n=26) and the "broad-sense natto group" (n=27) are indicated by orange lines. Key strains discussed in the text, the systematic nearest neighbor (Food_NPL_1) and the functional nearest neighbor (Food_IND_1), are highlighted with green squares on their labels.

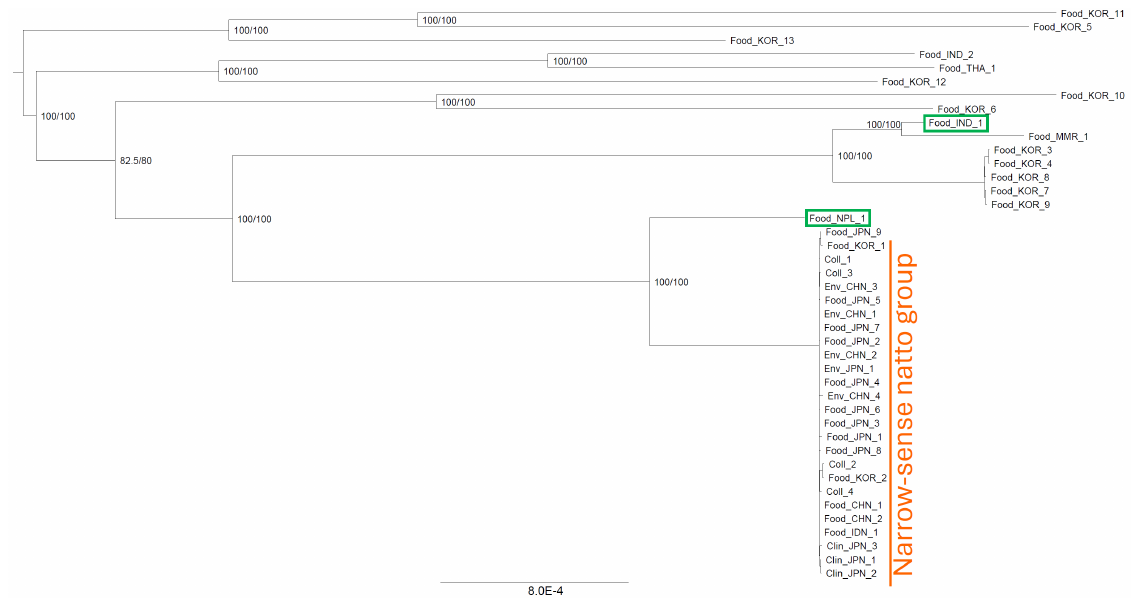

**Supplementary Figure 5.** Core-genome-based phylogenetic tree of the 42 *Bacillus subtilis* strains with support values. This phylogenetic tree was constructed from the core genome alignment of the 42 strains using the maximum-likelihood method with IQ-TREE 2. The tree is displayed using midpoint rooting. Numbers at the nodes indicate support values from SH-aLRT and UFBoot. Support values for tightly clustered strains are not shown for clarity. The "narrow-sense natto group" (n=26) is indicated by an orange line. Key strains discussed in the text, the systematic nearest neighbor (Food_NPL_1) and the functional nearest neighbor (Food_IND_1), are highlighted with green squares on their labels.

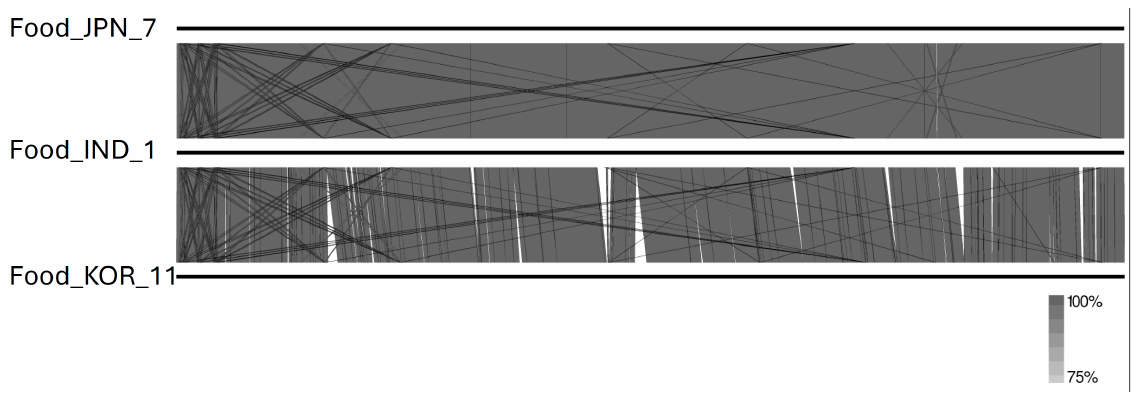

**Supplementary Figure 6.** Chromosome-wide synteny analysis of representative strains. This visualization compares the complete chromosome sequences of three strains, represented by horizontal bars: *B. subtilis* var. *natto* BEST195 (Food_JPN_7, top bar), the bekang isolate (Food_IND_1, middle bar), and *B. subtilis* KFRI-P74 (Food_KOR_11, bottom bar). The shaded regions connecting the genomes denote areas of nucleotide similarity identified by BLASTn, with the grayscale intensity corresponding to the percentage identity shown in the scale bar. The top panel shows the pairwise comparison between the natto strain (Food_JPN_7) and the bekang isolate (Food_IND_1). The bottom panel shows the pairwise comparison between the bekang isolate (Food_IND_1) and the "natto-related group" strain (Food_KOR_11).

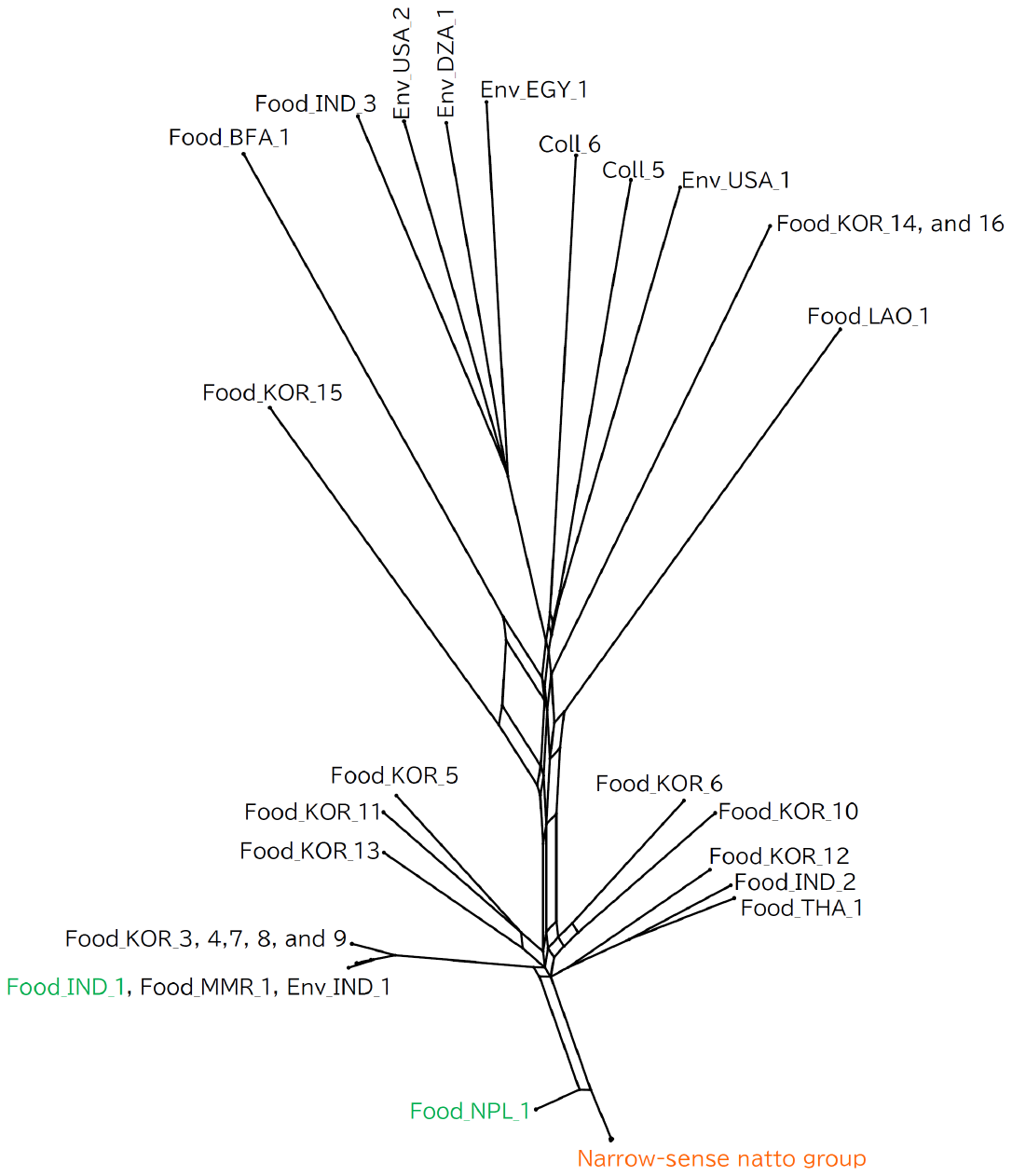

**Supplementary Figure 7.** Phylogenetic network of 55 *B. subtilis* strains. The network was constructed from the core genome alignment using the Neighbor-Net algorithm. The tree-like portions of the network depict vertical, clonal inheritance, while the reticulate (box-like) structures indicate conflicting phylogenetic signals suggestive of recombination or horizontal gene transfer events.

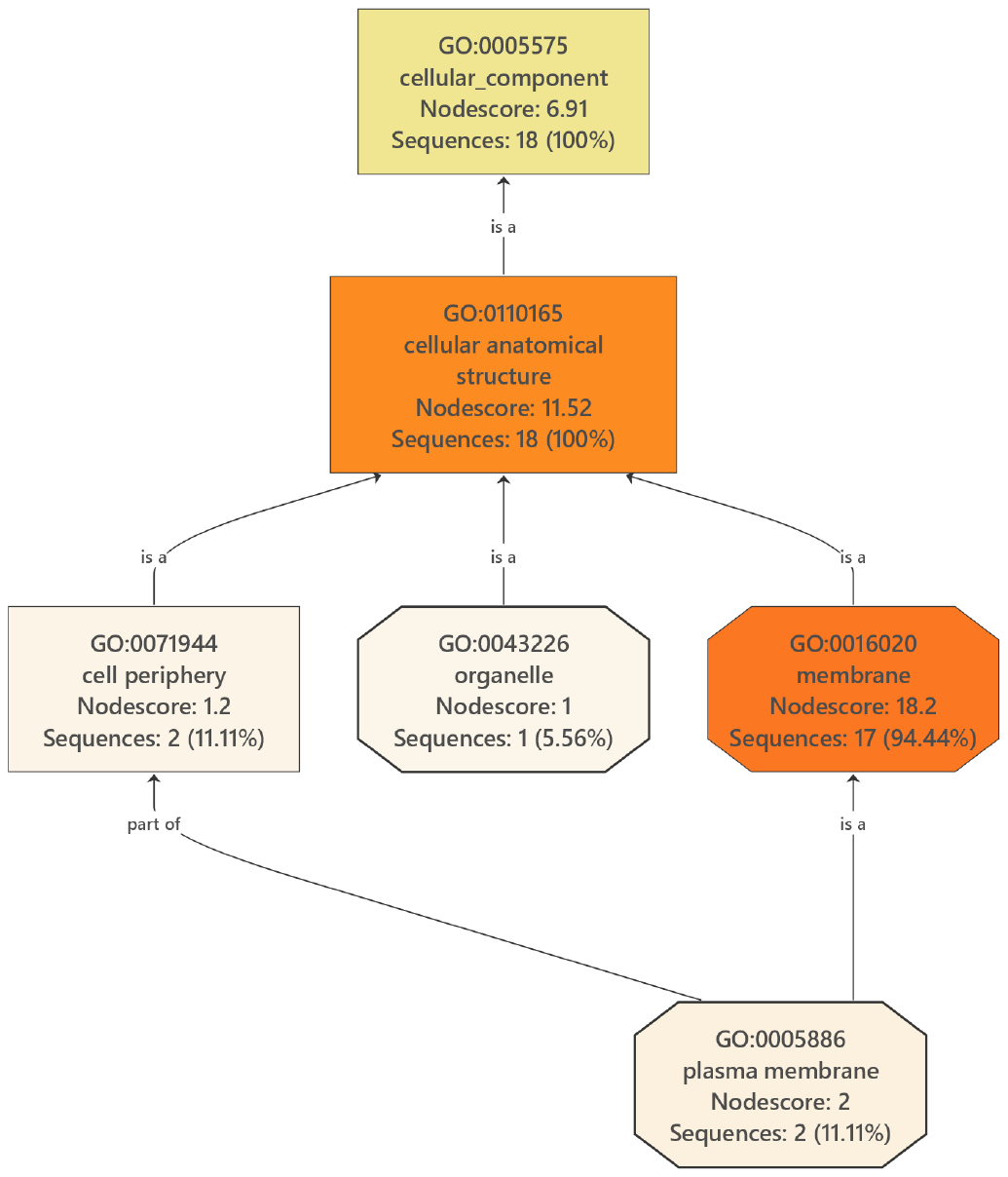
**Supplementary Figure 8.** Gene Ontology (GO) enrichment analysis for the Cellular Component (CC) category of the "broad-sense natto group"-specific gene set. The figure shows a Directed Acyclic Graph of enriched GO terms. Each box represents a GO term, displaying its ID, description, and the number of associated gene sequences. The color intensity of the boxes is proportional to the significance of enrichment, with more significant terms shown in shades of orange. The connecting lines represent hierarchical relationships between the terms.

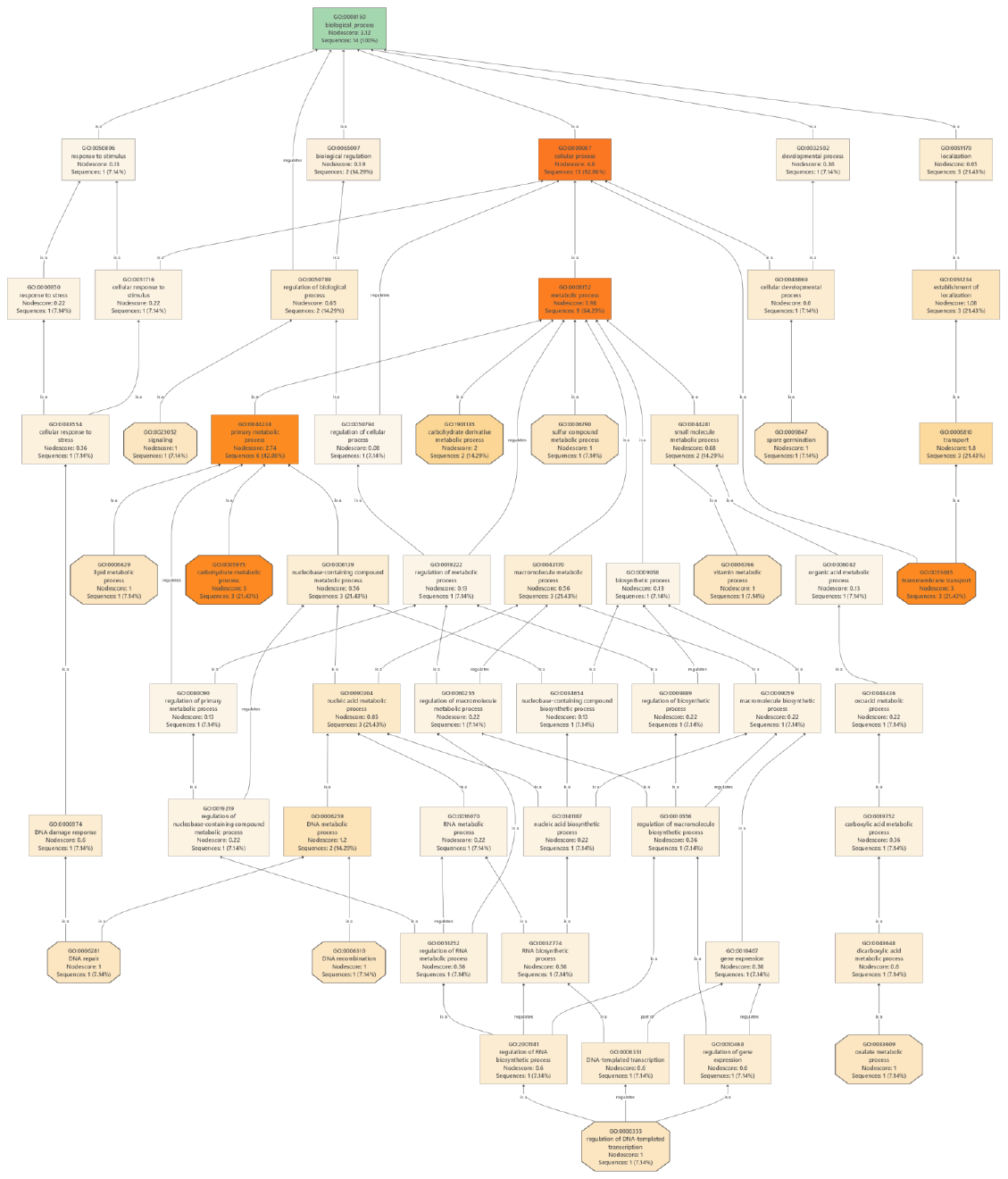

**Supplementary Figure 9.** Gene Ontology (GO) enrichment analysis for the Biological Process (BP) category of the "natto-related group"-specific gene set. The figure shows a Directed Acyclic Graph of enriched GO terms, generated using OmicsBox. Each node (box) represents a GO term and displays its ID, description, NodeScore, and the number of associated gene sequences. The color intensity of each node is proportional to its enrichment significance, with more significantly enriched terms shown in shades of orange. The connecting lines represent hierarchical relationships between the terms.

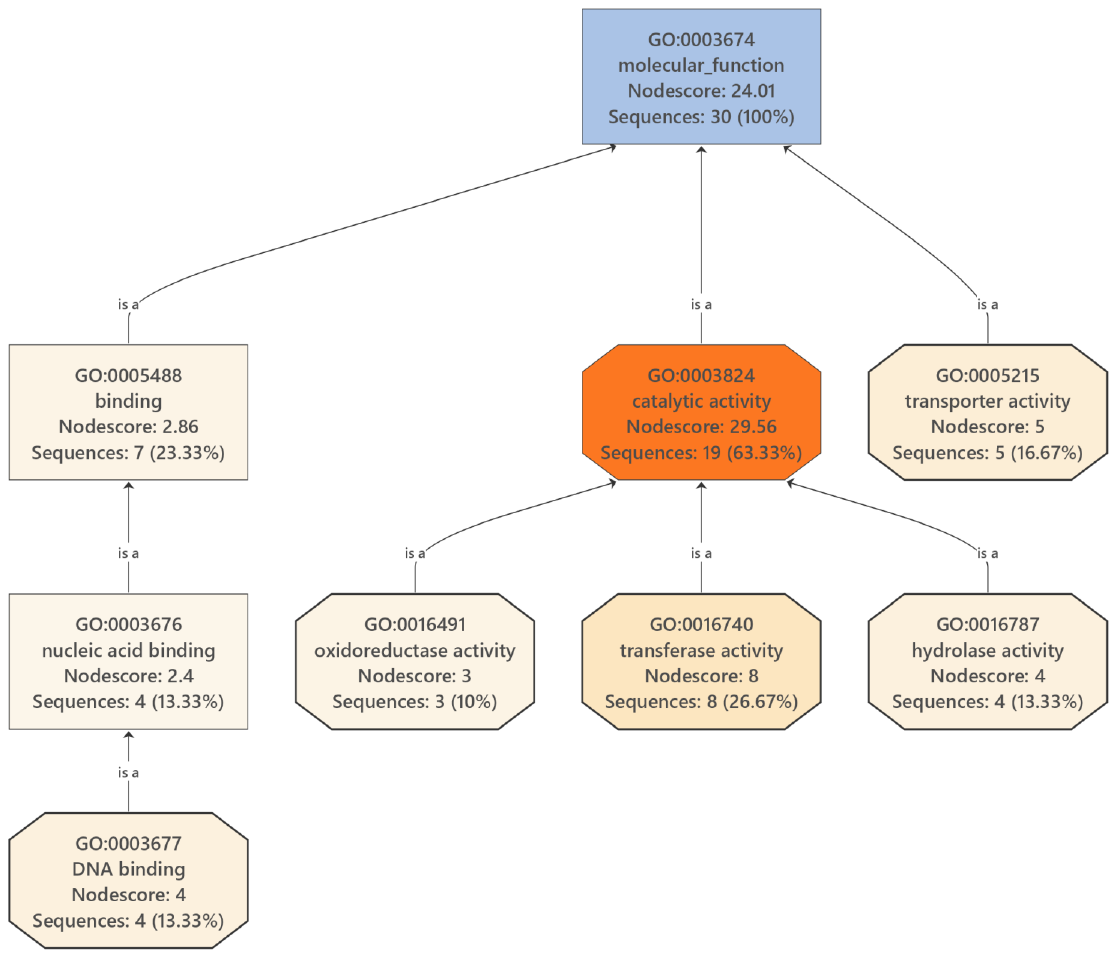

**Supplementary Figure 10.** Gene Ontology (GO) enrichment analysis for the Molecular Function (MF) category of the "natto-related group"-specific gene set. The figure shows a Directed Acyclic Graph of enriched GO terms. Each box represents a GO term, displaying its ID, description, and the number of associated gene sequences. The color intensity of the boxes is proportional to the significance of enrichment, with more significant terms shown in shades of orange. The connecting lines represent hierarchical relationships between the terms.

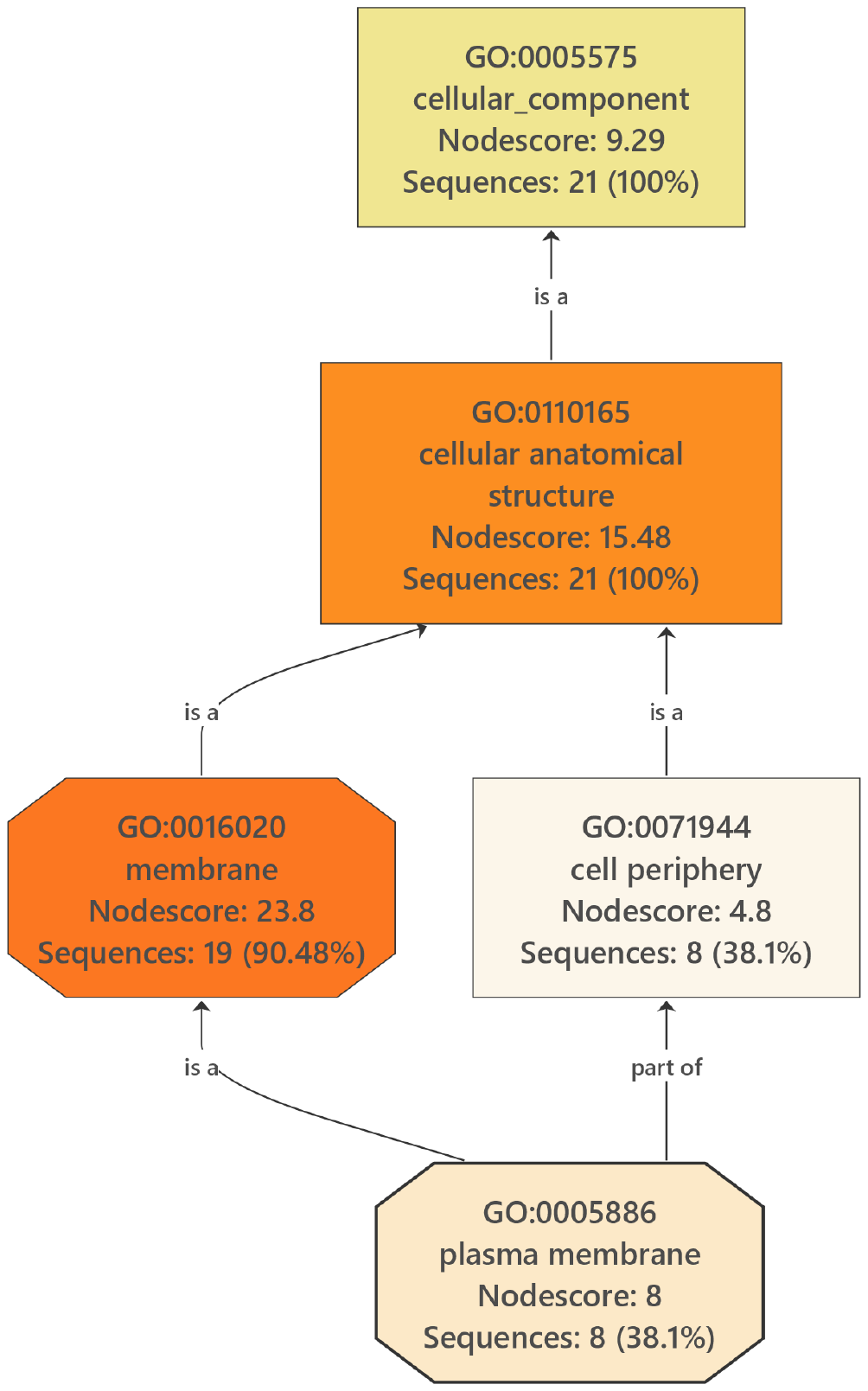

**Supplementary Figure 11.** Gene Ontology (GO) enrichment analysis for the Cellular Component (CC) category of the "natto-related group"-specific gene set. The figure shows a Directed Acyclic Graph of enriched GO terms. Each box represents a GO term, displaying its ID, description, and the number of associated gene sequences. The color intensity of the boxes is proportional to the significance of enrichment, with more significant terms shown in shades of orange. The connecting lines represent hierarchical relationships between the terms.

**Supplementary Table 1.** Genome assembly quality metrics for the 55 *Bacillus subtilis* strains used in this study.

| Name | Completeness | Contamination |
| --- | --- | --- |
| Clin_JPN_1 | 100 | 0.08 |
| Clin_JPN_2 | 100 | 0.07 |
| Clin_JPN_3 | 100 | 0.07 |
| Coll_1 | 100 | 0.09 |
| Coll_2 | 100 | 0.07 |
| Coll_3 | 100 | 0.05 |
| Coll_4 | 100 | 0.08 |
| Env_CHN_1 | 99.99 | 0.13 |
| Env_CHN_2 | 100 | 0.07 |
| Env_CHN_3 | 100 | 0.08 |
| Env_CHN_4 | 100 | 0.08 |
| Env_JPN_1 | 100 | 0.08 |
| Food_CHN_1 | 100 | 0.08 |
| Food_CHN_2 | 100 | 0.08 |
| Food_IDN_1 | 100 | 0.07 |
| Food_IND_1 | 100 | 0.06 |
| Food_JPN_1 | 100 | 0.11 |
| Food_JPN_2 | 100 | 0.07 |
| Food_JPN_3 | 100 | 0.08 |
| Food_JPN_4 | 100 | 0.08 |
| Food_JPN_5 | 100 | 0.08 |
| Food_JPN_6 | 100 | 0.08 |
| Food_JPN_7 | 100 | 0.09 |
| Food_JPN_8 | 100 | 0.08 |
| Food_JPN_9 | 100 | 0.07 |
| Food_KOR_1 | 100 | 0.23 |
| Food_KOR_2 | 100 | 0.07 |
| Food_IND_2 | 100 | 0.14 |
| Food_KOR_3 | 100 | 0.04 |
| Food_KOR_4 | 100 | 0.04 |
| Food_KOR_5 | 100 | 0.02 |
| Food_KOR_6 | 100 | 0 |
| Food_KOR_7 | 100 | 0.14 |
| Food_KOR_8 | 100 | 0.06 |
| Food_KOR_9 | 100 | 0.05 |
| Food_KOR_10 | 100 | 0.05 |
| Food_KOR_11 | 100 | 0.04 |
| Food_KOR_12 | 100 | 0.09 |
| Food_KOR_13 | 100 | 0.05 |
| Food_MMR_1 | 100 | 0.03 |
| Food_NPL_1 | 100 | 0.08 |
| Food_THA_1 | 100 | 0.17 |
| Coll_5 | 100 | 0.05 |
| Coll_6 | 100 | 0.09 |
| Env_DZA_1 | 100 | 0.02 |
| Env_EGY_1 | 100 | 0.06 |
| Env_IND_1 | 100 | 0.04 |
| Env_USA_1 | 100 | 0.01 |
| Env_USA_2 | 100 | 0.02 |
| Food_BFA_1 | 100 | 0.22 |
| Food_IND_3 | 100 | 0.02 |
| Food_KOR_14 | 100 | 0.64 |
| Food_KOR_15 | 100 | 0.05 |
| Food_KOR_16 | 100 | 0.87 |
| Food_LAO_1 | 100 | 0.29 |
