## Supplementary material for "Conserved accessory genes link a phylogenetically distinct *Bacillus subtilis* strain from Indian bekang to the Japanese natto clade": Dataset 13: analyze_genes_B_specific.py

```
#!/usr/bin/env python3

import pandas as pd
import numpy as np
import argparse

def analyze_genes_B_specific(input_file, output_file):
    """
    Analyzes gene presence/absence data, compares group A and group B to extract
    genes specific to group B, and outputs the results to a CSV file.
    The 'ratio' column is calculated as (count in Group A / count in Group B) * 100,
    and the table is sorted by the 'ratio' column in ascending order.
    Header names are changed to specified ones.

    Args:
        input_file (str): Path to the input CSV file.
        output_file (str): Path to the output CSV file.
    """
    try:
        df = pd.read_csv(input_file)
        print(f"Successfully loaded '{input_file}'.")
    except FileNotFoundError:
        print(f"Error: '{input_file}' not found. Please check the file path.")
        return
    except Exception as e:
        print(f"An error occurred while reading the file: {e}")
        return

    # Set the first column as the index (gene name)
    df = df.set_index(df.columns[0])

    # Identify the columns for each group
    cols_A = [col for col in df.columns if col.startswith('A_')]
    cols_B = [col for col in df.columns if col.startswith('B_')]

    if not cols_A or not cols_B:
        print("Error: Columns starting with 'A_' or 'B_' were not found. Please check the column names.")
        return

    print(f"Found {len(cols_A)} strains in Group A.")
    print(f"Found {len(cols_B)} strains in Group B.")

    # Binarize the data: 1 for non-missing (present), 0 for missing (absent)
    df_numeric = df.notna().astype(int)

    # Count the number of strains where each gene is present in each group
    df['count_A'] = df_numeric[cols_A].sum(axis=1)
    df['count_B'] = df_numeric[cols_B].sum(axis=1)

    num_A = len(cols_A)
    num_B = len(cols_B)

    # Calculate the percentage of strains where each gene is present in each group
    df['perc_A'] = df['count_A'] / num_A
    df['perc_B'] = df['count_B'] / num_B

    # Filter 1: Extract genes present in >= 80% of Group B
    df_filtered1 = df[df['perc_B'] >= 0.8].copy()
    print(f"Number of genes after Filter 1 (B >= 80%): {len(df_filtered1)}")

    # Filter 2: Extract genes present in <= 20% of Group A
    df_final = df_filtered1[df_filtered1['perc_A'] <= 0.2].copy()
    print(f"Number of genes after Filter 2 (A <= 20%): {len(df_final)}")

    # Create a DataFrame for the results
    df_result = pd.DataFrame(index=df_final.index)
    df_result['count_B_in_filtered'] = df_final['count_B'] # Number of strains included in group B
    df_result['count_A_in_filtered'] = df_final['count_A'] # Number of strains included in group A
    
    # Calculate ratio: (count in Group A / count in Group B) * 100
    # Avoid division by zero by replacing 0 with NaN for the calculation
    df_result['ratio_A_div_B'] = (df_result['count_A_in_filtered'] / df_result['count_B_in_filtered'].replace(0, np.nan)) * 100
    
    # Join the original data columns (only those starting with 'A_' or 'B_')
    original_data_cols_AB = [col for col in df.columns if col.startswith(('A_', 'B_'))]
    df_result = df_result.join(df_final[original_data_cols_AB]) # Join from df_final

    df_result = df_result.reset_index()
    # Rename the first column (gene name) to 'gene'
    df_result = df_result.rename(columns={df_result.columns[0]: 'gene'})

    # Specify the column order
    # Place gene, count_B_in_filtered, count_A_in_filtered, ratio_A_div_B at the beginning, followed by the original data columns
    cols_order = ['gene', 'count_B_in_filtered', 'count_A_in_filtered', 'ratio_A_div_B'] + original_data_cols_AB
    df_result = df_result[cols_order]
    
    # Sort by the 'ratio_A_div_B' column in ascending order
    df_result.sort_values(by='ratio_A_div_B', ascending=True, inplace=True)

    # Rename columns to the specified names
    df_result.rename(columns={
        'count_B_in_filtered': 'The number included in group B',
        'count_A_in_filtered': 'The number included in group A',
        'ratio_A_div_B': 'ratio'
    }, inplace=True)

    try:
        df_result.to_csv(output_file, index=False)
        print(f"\nAnalysis complete. Results have been saved to '{output_file}'.")
        print(f"The final table contains {len(df_result)} genes.")
    except Exception as e:
        print(f"An error occurred while writing the file: {e}")

if __name__ == "__main__":
    parser = argparse.ArgumentParser(description='Analyzes bacterial gene presence/absence data to compare Group A and Group B and extract genes specific to Group B.')
    parser.add_argument('-i', '--input', required=True, help='Input CSV file name')
    parser.add_argument('-o', '--output', required=True, help='Output CSV file name')
    args = parser.parse_args()

    analyze_genes_B_specific(args.input, args.output)
```
