## Supplementary material for "Conserved accessory genes link a phylogenetically distinct *Bacillus subtilis* strain from Indian bekang to the Japanese natto clade": Dataset 13: process_csv.py

```
#!/usr/bin/env python3

import csv
import argparse
import sys

def process_csv(input_file):
    """
    Processes a CSV file and outputs it to standard output in a specified format.

    Args:
        input_file (str): The path to the input CSV file.
    """
    try:
        with open(input_file, 'r', newline='', encoding='utf-8') as infile:
            reader = csv.reader(infile)

            # Skip the header row
            try:
                next(reader)
            except StopIteration:
                # Do nothing if the file is empty or contains only a header
                return

            # Start processing from the second row
            for row in reader:
                if not row:  # Skip empty rows
                    continue

                # Extract the first column
                col1 = row[0] if len(row) > 0 else ""

                # Search from the 6th column onwards
                found_value = ""
                if len(row) > 5 and row[5]:
                    found_value = row[5]
                else:
                    # If the 6th column is empty, search from the 7th column onwards
                    for i in range(6, len(row)):
                        if row[i]:
                            found_value = row[i]
                            break

                # Print the result to standard output, separated by a tab
                print(f"{col1}\t{found_value}")

    except FileNotFoundError:
        print(f"Error: File '{input_file}' not found.", file=sys.stderr)
    except Exception as e:
        print(f"Error: {e}", file=sys.stderr)

def main():
    """
    Parses command-line arguments and calls the CSV processing function.
    """
    parser = argparse.ArgumentParser(description="Processes a CSV file, extracts specified columns, and prints to standard output.")
    parser.add_argument("-i", "--input", required=True, help="Path to the input CSV file.")

    args = parser.parse_args()
    process_csv(args.input)

if __name__ == "__main__":
    main()
```
