## Supplementary material for "Conserved accessory genes link a phylogenetically distinct *Bacillus subtilis* strain from Indian bekang to the Japanese natto clade": Dataset 13: process_fasta.py

```
#!/usr/bin/env python3
# -*- coding: utf-8 -*-

import argparse
import os
import sys

def parse_fasta(file_path):
    """
    Parses the specified FASTA file and returns a dictionary with the
    header (first word) as the key and the sequence as the value.

    Args:
        file_path (str): The path to the FASTA file.

    Returns:
        dict: A dictionary in the format {header: sequence}.
    """
    sequences = {}
    current_header = None
    with open(file_path, 'r') as f:
        for line in f:
            line = line.strip()
            if not line:
                continue
            if line.startswith('>'):
                # Use the first word after '>' as the header (gene name).
                current_header = line[1:].split()[0]
                sequences[current_header] = ""
            elif current_header:
                sequences[current_header] += line
    return sequences

def read_mapping(file_path):
    """
    Reads a tab-separated mapping file and returns a dictionary
    mapping gene names from FASTA to output gene names, and a list of
    FASTA gene names in the order of the mapping file.

    Args:
        file_path (str): The path to the mapping file.

    Returns:
        tuple: (
            dict: {fasta_gene_name: output_gene_name},
            list: [fasta_gene_name, ...]
        )
    """
    mapping = {}
    ordered_names = []
    try:
        with open(file_path, 'r') as f:
            for line_num, line in enumerate(f, 1):
                line = line.strip()
                if not line or line.startswith('#'):
                    continue
                parts = line.split('\t')
                if len(parts) >= 2:
                    output_name = parts[0]
                    fasta_name = parts[1]
                    mapping[fasta_name] = output_name
                    ordered_names.append(fasta_name)
                else:
                    print(f"Warning: Line {line_num} in the mapping file has an invalid format: {line}", file=sys.stderr)
    except FileNotFoundError:
        print(f"Error: Mapping file '{file_path}' not found.", file=sys.stderr)
        sys.exit(1)
    return mapping, ordered_names

def write_output_fasta(output_handle, ordered_fasta_names, fasta_to_output_map, all_sequences):
    """
    Writes sequences in FASTA format to the specified output handle.

    Args:
        output_handle: The file handle to write to (e.g., sys.stdout or a file opened with open()).
        ordered_fasta_names (list): A list of FASTA gene names in the desired output order.
        fasta_to_output_map (dict): A mapping from FASTA names to output names.
        all_sequences (dict): All sequence data.
    """
    print("Info: Generating output FASTA file...", file=sys.stderr)
    found_count = 0
    missing_count = 0

    for fasta_name in ordered_fasta_names:
        if fasta_name in all_sequences:
            output_name = fasta_to_output_map[fasta_name]
            sequence = all_sequences[fasta_name]
            print(f">{output_name}", file=output_handle)
            # Wrap the sequence to 60 characters per line for output
            for i in range(0, len(sequence), 60):
                print(sequence[i:i+60], file=output_handle)
            found_count += 1
        else:
            output_name = fasta_to_output_map.get(fasta_name, "Unknown")
            print(f"Warning: Mapped gene '{fasta_name}' (output name: '{output_name}') was not found in the FASTA files.", file=sys.stderr)
            missing_count += 1

    print(f"Info: Processing complete. {found_count} genes were written, and {missing_count} were not found.", file=sys.stderr)


def main():
    """
    Main function. Parses arguments, processes FASTA files, and generates the output.
    """
    parser = argparse.ArgumentParser(
        description="Generates a FASTA file with specified gene names from multiple FASTA files in a directory and a mapping file."
    )
    parser.add_argument(
        '-I', '--input_dir',
        required=True,
        help="Path to the directory containing multiple FASTA files."
    )
    parser.add_argument(
        '-i', '--mapping_file',
        required=True,
        help="Path to the tab-separated mapping file (Column 1: output gene name, Column 2: gene name in FASTA)."
    )
    parser.add_argument(
        '-o', '--output_file',
        help="Path for the output FASTA file. If not specified, output will be written to stdout."
    )
    args = parser.parse_args()

    # Read the mapping file
    fasta_to_output_map, ordered_fasta_names = read_mapping(args.mapping_file)

    # Check if the input directory exists
    if not os.path.isdir(args.input_dir):
        print(f"Error: Directory '{args.input_dir}' not found.", file=sys.stderr)
        sys.exit(1)

    # Read all FASTA files in the directory
    all_sequences = {}
    print(f"Info: Reading FASTA files from directory '{args.input_dir}'...", file=sys.stderr)
    for filename in os.listdir(args.input_dir):
        file_path = os.path.join(args.input_dir, filename)
        if os.path.isfile(file_path):
            print(f"  - Processing file '{filename}'...", file=sys.stderr)
            try:
                sequences_in_file = parse_fasta(file_path)
                all_sequences.update(sequences_in_file)
            except Exception as e:
                print(f"Warning: An error occurred while processing file '{filename}': {e}", file=sys.stderr)
    print(f"Info: Loaded {len(all_sequences)} sequences.", file=sys.stderr)

    # Write output in FASTA format based on the mapping file
    if args.output_file:
        # If -o is specified: write to a file
        print(f"Info: Output will be written to file '{args.output_file}'.", file=sys.stderr)
        try:
            with open(args.output_file, 'w') as output_handle:
                write_output_fasta(output_handle, ordered_fasta_names, fasta_to_output_map, all_sequences)
        except IOError as e:
            print(f"Error: Could not open output file '{args.output_file}': {e}", file=sys.stderr)
            sys.exit(1)
    else:
        # If -o is not specified: write to standard output
        print("Info: Output will be written to standard output.", file=sys.stderr)
        write_output_fasta(sys.stdout, ordered_fasta_names, fasta_to_output_map, all_sequences)


if __name__ == "__main__":
    main()
```
