## Supplementary material for "Conserved accessory genes link a phylogenetically distinct *Bacillus subtilis* strain from Indian bekang to the Japanese natto clade": Dataset 13: roary_plots_2.py

```
#!/usr/bin/env python
# Copyright (C) <2015> EMBL-European Bioinformatics Institute

# This program is free software: you can redistribute it and/or
# modify it under the terms of the GNU General Public License as
# published by the Free Software Foundation, either version 3 of
# the License, or (at your option) any later version.

# This program is distributed in the hope that it will be useful,
# but WITHOUT ANY WARRANTY; without even the implied warranty of
# MERCHANTABILITY or FITNESS FOR A PARTICULAR PURPOSE. See the
# GNU General Public License for more details.

# Neither the institution name nor the name roary_plots
# can be used to endorse or promote products derived from
# this software without prior written permission.
# For written permission, please contact <>.

# Products derived from this software may not be called roary_plots
# nor may roary_plots appear in their names without prior written
# permission of the developers. You should have received a copy
# of the GNU General Public License along with this program.
# If not, see <http://www.gnu.org/licenses/>.

__author__ = "Marco Galardini"
__version__ = '0.1.0'

def get_options():
    import argparse

    # create the top-level parser
    description = "Create plots from roary outputs"
    parser = argparse.ArgumentParser(description = description,
                                     prog = 'roary_plots.py')

    parser.add_argument('tree', action='store',
                        help='Newick Tree file', default='accessory_binary_genes.fa.newick')
    parser.add_argument('spreadsheet', action='store',
                        help='Roary gene presence/absence spreadsheet', default='gene_presence_absence.csv')

    parser.add_argument('--labels', action='store_true',
                        default=False,
                        help='Add node labels to the tree (up to 10 chars)')
    parser.add_argument('--format',
                        choices=('png',
                                 'tiff',
                                 'pdf',
                                 'svg'),
                        default='png',
                        help='Output format [Default: png]')
    parser.add_argument('-N', '--skipped-columns', action='store',
                        type=int,
                        default=14,
                        help='First N columns of Roary\'s output to exclude [Default: 14]')

    parser.add_argument('--version', action='version',
                        version='%(prog)s '+__version__)

    return parser.parse_args()

if __name__ == "__main__":
    options = get_options()

    import matplotlib
    matplotlib.use('Agg') # Use Agg backend for non-interactive plotting

    import matplotlib.pyplot as plt
    import seaborn as sns

    sns.set_style('white')

    import os
    import pandas as pd
    import numpy as np
    from Bio import Phylo

    # Load the tree
    t = Phylo.read(options.tree, 'newick')

    # --- MODIFICATION START ---
    # Root the tree at midpoint
    t.root_at_midpoint()
    # --- MODIFICATION END ---

    # Max distance to create better plots (recalculate after rooting)
    mdist = max([t.distance(t.root, x) for x in t.get_terminals()])
    if mdist == 0: # Handle cases where all branch lengths are zero or tree has one terminal
        mdist = 1 

    # Load roary
    roary = pd.read_csv(options.spreadsheet, low_memory=False)
    # Set index (group name)
    roary.set_index('Gene', inplace=True)
    # Drop the other info columns
    roary.drop(list(roary.columns[:options.skipped_columns-1]), axis=1, inplace=True)

    # Transform it in a presence/absence matrix (1/0)
    roary.replace('.{2,100}', 1, regex=True, inplace=True)
    roary.replace(np.nan, 0, regex=True, inplace=True) # Ensure NaN is replaced with 0

    # Sort the matrix by the sum of strains presence
    idx = roary.sum(axis=1).sort_values(ascending=False).index
    roary_sorted = roary.loc[idx]

    # Pangenome frequency plot
    plt.figure(figsize=(7, 5))

    plt.hist(roary.sum(axis=1), bins=min(roary.shape[1], 50), # Added bins for clarity
             histtype="stepfilled", alpha=.7)

    plt.xlabel('No. of genomes')
    plt.ylabel('No. of genes')

    sns.despine(left=True,
                bottom=True)
    plt.savefig('pangenome_frequency.%s'%options.format, dpi=300)
    plt.clf() # Clear the current figure

    # Sort the matrix according to tip labels in the tree
    # Ensure all terminal names are strings for proper indexing
    terminal_names = [str(x.name) for x in t.get_terminals()]
    # Filter roary_sorted columns to only include those present in the tree
    roary_sorted = roary_sorted[[name for name in terminal_names if name in roary_sorted.columns]]
    # Reorder terminal_names to match the filtered roary_sorted columns
    # This is important if some tree labels are not in the roary matrix
    ordered_terminal_names = [name for name in terminal_names if name in roary_sorted.columns]


    # Plot presence/absence matrix against the tree
    with sns.axes_style('whitegrid'):
        fig = plt.figure(figsize=(17, 10))

        ax1=plt.subplot2grid((1,40), (0, 10), colspan=30)
        # Ensure roary_sorted is reordered according to the tree's terminal order
        # after potential filtering
        # The matrix should be ordered by 'ordered_terminal_names'
        # And we transpose it for the heatmap
        matrix_to_plot = roary_sorted[ordered_terminal_names].T

        a=ax1.matshow(matrix_to_plot, cmap=plt.cm.Blues,
                      vmin=0, vmax=1,
                      aspect='auto',
                      interpolation='none',
                      )
        ax1.set_yticks([])
        ax1.set_xticks([])
        ax1.axis('off')

        # ax = fig.add_subplot(1,2,1) # This line seems problematic with subplot2grid
        # matplotlib v1/2 workaround
        try:
            ax=plt.subplot2grid((1,40), (0, 0), colspan=10, facecolor='white')
        except AttributeError: # older matplotlib
            ax=plt.subplot2grid((1,40), (0, 0), colspan=10, axisbg='white')

        fig.subplots_adjust(wspace=0, hspace=0)

        ax1.set_title('Roary matrix\n(%d gene clusters)'%roary.shape[0])

        # Adjust xlim for the tree plot, especially if labels are shown
        # The xlim calculation for labels needs careful adjustment
        # Ensure mdist is positive to avoid issues with xlim
        if mdist <= 0:
            effective_mdist = 1.0 # Default positive value if mdist is not positive
        else:
            effective_mdist = mdist

        xlim_right_no_labels = effective_mdist + effective_mdist * 0.1
        # Adjust xlim if labels are present
        # The original xlim for labels was: mdist+mdist*0.45-mdist*roary.shape[1]*0.001
        # This can become negative or too small if roary.shape[1] (number of strains) is large.
        # We need a more robust calculation for xlim when labels are shown.
        # Let's make the label space proportional to mdist but ensure it's reasonable.
        xlim_right_labels = effective_mdist + effective_mdist * 0.6 # Increased space for labels

        if options.labels:
            fsize = 12 - 0.1*len(ordered_terminal_names) # Use actual number of plotted strains
            if fsize < 3:
                fsize = 3
            with plt.rc_context({'font.size': fsize}):
                Phylo.draw(t, axes=ax,
                           show_confidence=False,
                           label_func=lambda x: str(x.name)[:40] if x.is_terminal() else None, # Show labels only for terminals
                           xticks=([],), yticks=([],),
                           ylabel=('',), xlabel=('',),
                           # xlim ensures tree is not cut off and labels have space
                           xlim=(-effective_mdist*0.1, xlim_right_labels),
                           axis=('off',),
                           title=('Tree\n(%d strains)'%len(ordered_terminal_names),), # Use actual number of plotted strains
                           do_show=False,
                           label_colors=lambda x: 'black' # Ensure labels are visible
                          )
        else:
            Phylo.draw(t, axes=ax,
                       show_confidence=False,
                       label_func=lambda x: None,
                       xticks=([],), yticks=([],),
                       ylabel=('',), xlabel=('',),
                       xlim=(-effective_mdist*0.1, xlim_right_no_labels),
                       axis=('off',),
                       title=('Tree\n(%d strains)'%len(ordered_terminal_names),), # Use actual number of plotted strains
                       do_show=False,
                      )
        plt.savefig('pangenome_matrix.%s'%options.format, dpi=300)
        plt.clf() # Clear the current figure

    # Plot the pangenome pie chart
    plt.figure(figsize=(10, 10)) # New figure for pie chart

    num_genomes = roary.shape[1] # Number of genomes in the matrix

    core     = roary[(roary.sum(axis=1) >= num_genomes*0.99) & (roary.sum(axis=1) <= num_genomes)].shape[0]
    softcore = roary[(roary.sum(axis=1) >= num_genomes*0.95) & (roary.sum(axis=1) <  num_genomes*0.99)].shape[0]
    shell    = roary[(roary.sum(axis=1) >= num_genomes*0.15) & (roary.sum(axis=1) <  num_genomes*0.95)].shape[0]
    cloud    = roary[roary.sum(axis=1)  < num_genomes*0.15].shape[0]

    total = roary.shape[0]

    def my_autopct(pct):
        if total == 0: # Avoid division by zero if roary matrix is empty
            return '0'
        val=int(round(pct*total/100.0))
        return '{v:d}'.format(v=val)

    # Ensure slices are non-negative
    slices = [max(0, core), max(0, softcore), max(0, shell), max(0, cloud)]
    labels_pie = [
        'core\n(%.0f%% <= strains <= 100%%)' % (0.99 * 100),
        'soft-core\n(%.0f%% <= strains < %.0f%%)' % (0.95 * 100, 0.99 * 100),
        'shell\n(%.0f%% <= strains < %.0f%%)' % (0.15 * 100, 0.95 * 100),
        'cloud\n(strains < %.0f%%)' % (0.15 * 100)
    ]
    
    # Filter out zero-value slices and corresponding labels to avoid warnings/errors
    non_zero_slices = [s for s in slices if s > 0]
    non_zero_labels = [labels_pie[i] for i, s in enumerate(slices) if s > 0]
    non_zero_explode = [([0.1, 0.05, 0.02, 0])[i] for i, s in enumerate(slices) if s > 0] # Adjust explode accordingly
    
    # Colors based on the original logic, ensure division by total is safe
    if total > 0:
        colors_pie = [(0, 0, 1, float(x)/total) if total > 0 else (0,0,1,0) for x in slices]
        non_zero_colors = [colors_pie[i] for i,s in enumerate(slices) if s > 0]
    else: # Default colors if total is 0
        non_zero_colors = ['blue', 'lightblue', 'gray', 'lightgray'][:len(non_zero_slices)]


    if total > 0 and sum(non_zero_slices) > 0: # Only plot if there's data
        plt.pie(non_zero_slices,
            labels=non_zero_labels,
            explode=non_zero_explode, radius=0.9,
            colors=non_zero_colors,
            autopct=my_autopct)
        plt.title('Pangenome distribution (%d gene clusters total)' % total, pad=20)
    else:
        plt.text(0.5, 0.5, "No data to display in pie chart", ha='center', va='center')
        plt.title('Pangenome distribution (0 gene clusters total)', pad=20)

    plt.savefig('pangenome_pie.%s'%options.format, dpi=300)
    plt.clf() # Clear the current figure

    print("Roary plots generated successfully.")
```
